# LRRK2 I1371V Impairs Astrocytic Glucose Metabolism and Triggers Multi-Organellar Dysfunction in PD

**DOI:** 10.64898/2026.01.19.700314

**Authors:** Roon Banerjee, Rashmi Santhoshkumar, Vikram Holla, Nitish Kamble, Ravi Yadav, Pramod Kumar Pal, Indrani Datta

## Abstract

While LRRK2 mutations modulate systemic glucose homeostasis and metabolic dysfunction precedes Parkinson’s disease (PD) motor-symptoms, the impact of pathogenic LRRK2-mutations on astrocytic glucose-uptake and organellar function remains unexplored. Here, we demonstrate that LRRK2-I1371V mutation causes profound metabolic and organellar dysfunction in LRRK2-I1371V PD-iPSC-derived astrocytes and U87 cells overexpressing I1371V variant. LRRK2-I1371V astrocytes exhibit significantly reduced GLUT1 expression and plasma membrane localization, resulting in impaired glucose-uptake and decreased lactate-production. This metabolic insufficiency correlates with cascading mitochondrial dysfunction, characterized by membrane depolarization, elevated reactive-oxygen-species, enhanced ubiquitination and reduced proteasomal-activity. Reduced LAMP1/LAMP2 expression, impaired lysosomal-acidification, and selective cathepsin D deficiency were observed. Accumulation of undegraded cargo was confirmed by transmission-electron-microscopy upon α-synuclein exposure. ER stress was evident through upregulated GADD34/CHOP, increased phospho-PERK, and reduced nascent protein synthesis. Our results reveal that LRRK2-I1371V induces glucose-uptake deficits, causing energy depletion and multi-organellar dysfunction, identifying astrocytic metabolic restoration as a promising therapeutic target for I1371V-associated PD.

## Introduction

Parkinson’s disease (PD) is a progressive neurodegenerative disorder characterized by the selective degeneration of dopaminergic neurons (Ramesh et al., 2023), the accumulation of α-synuclein aggregates (Agarwal et al., 2022), and widespread glial dysfunction (Jeon et al., 2020; Domingues et al., 2020). Among the known genetic contributors, mutations in leucine-rich repeat kinase 2 (LRRK2) account for approximately 5–13% of familial and 1–5% of sporadic PD cases (Kumari et al., 2009). Similar to idiopathic PD, brains from LRRK2-mutant PD subjects exhibit significant dopaminergic neurodegeneration in the substantia nigra accompanied by α-synuclein-positive Lewy body pathology (Giasson et al. 2006, Ross et al. 2006).

LRRK2 encodes a large, multi-domain protein containing both kinase and GTPase activities, positioning it as a critical regulatory hub in multiple cellular processes including vesicle trafficking, autophagy, lysosomal function, and cytoskeletal dynamics (Nguyen et al. 2025). The protein’s complex architecture includes leucine-rich repeats, Ras-of-complex (Roc) GTPase domain, C-terminal of Roc (COR) linker region, kinase domain, and WD40 repeats, enabling diverse protein-protein interactions and enzymatic functions (Myasnikov et al. 2021).

Pathogenic mutations cluster primarily within the Roc-COR-kinase region. The most common pathogenic mutation is G2019S in the kinase domain, which enhances kinase activity (Greggio et al., 2009). Key mutations within the GTPase (ROC–COR) domain include R1441C/G/H, which alter GTPase function (Chang et al., 2017; Ali et al., 2022), and I1371V, located in the Roc domain (Paisán-Ruíz et al., 2005; Pankratz et al., 2006; Deng et al., 2008; Pirkevi et al., 2009; Seki et al., 2011; Sadhukhan et al., 2012; Janković et al., 2015; Lin et al., 2019; Fernández-Pajarín et al., 2023). Although GTPase variants produce classical PD symptoms, their clinical manifestations often differ from those associated with kinase domain mutations (Giordana et al., 2007; Cheon et al., 2012; Kalia et al., 2015; Huang et al., 2022; Taymans et al., 2023).

Current evidence suggests that LRRK2 mutations exert gain-of-function effects and contribute to neurodegeneration through multiple convergent mechanisms, including enhanced kinase activity, disrupted protein interactions, impaired vesicular trafficking, defective autophagy-lysosomal clearance, and altered synaptic function (Cookson et al. 2015; Boecker et al. 2023; Martin et al. 2014; Nikonova et al. 2012; Xiong et al. 2010).

The traditional neuron-centric view of PD has evolved to recognize the critical contributions of glial cells, particularly astrocytes, to disease pathogenesis and progression (Sonninen et al. 2020). While LRRK2 has been extensively studied in dopaminergic neurons, it is also expressed in astrocytes (Miklossy et al.2006; Booth et al. 2017). As the most abundant glial cell type in the brain, astrocytes play essential roles in maintaining neuronal health through antioxidant defense, metabolic support, neurotransmitter recycling, and α-synuclein clearance (Banker et al., 1980; Catalani et al., 2002; Perea et al., 2009; Henneberger et al., 2010; Bosson et al., 2015; Schreiner et al., 2015; van Deijk et al., 2017).

Studies on LRRK2 mutation-carrying astrocytes have reported impaired glutamate uptake and metabolism (Banerjee et al., 2022; Iovino et al., 2022), compromised Nrf2-mediated antioxidant systems (Banerjee et al. 2023), reduced levels of nerve growth factor, and increased levels of proinflammatory cytokines (interleukin-1β and tumor necrosis factor α) (Ho et al., 2024) and impaired ATP generation (Banerjee et al., 2023). However, regarding glucose uptake and metabolism, only a single study (Sonninen et al., 2020) in LRRK2 G2019S mutation-carrying astrocytes reported no significant differences in glucose transporter expression and glucose uptake.

The brain’s extraordinarily high energy demands, consuming approximately 20% of the body’s glucose despite representing only 2% of body weight, necessitate tight coupling between glucose availability and cellular function (Zhang et al. 2024). Astrocytes play a pivotal role in brain energy metabolism through glucose uptake, glycolytic processing, and lactate provision to neurons via the astrocyte-neuron lactate shuttle (ANLS)—a mechanism that supports synaptic transmission and neuronal survival, particularly during periods of high energy demand (Beard et al. 2022; Erlichman et al. 2008). This metabolic partnership is especially critical in the aging brain and during neurodegenerative processes, where energy deficits can precipitate cellular dysfunction and death (Kim et al. 2025).

Accumulating evidence indicates that metabolic dysfunction represents a core feature of PD. Neuroimaging studies reveal reduced glucose metabolism in affected brain regions prior to overt motor symptoms, and post-mortem analyses demonstrate mitochondrial respiratory chain defects and glycolytic enzyme alterations (Chen et al. 2025; Magistretti et al. 2015 Chen et al. 2023). Epidemiological studies linking diabetes and metabolic syndrome to increased PD risk further support the importance of glucose homeostasis in disease pathogenesis (Souza et al. 2021). Indeed, reduced expression of glucose transporters like GLUT1 limits neuronal glucose uptake, leading to ATP deficits, heightened oxidative stress, and ultimately acceleration of disease progression (Singh et al. 2025).

LRRK2 is expressed in several insulin-sensitive tissues, including muscle and adipose tissue, where it modulates insulin signalling and systemic glucose homeostasis [Kawakami et al. 2023; Imai et al. 2020; Funk et al. 2019]. LRRK2 is also expressed in the endocrine pancreas [Dule et al. 2025]. Its expression and the phosphorylation of its substrates Rab8a and Rab10 are markedly increased following high-fat diet treatment in wild-type mice (Kawakami et al. 2023). In cell cultures, pharmacological inhibition of LRRK2 enhances insulin-dependent GLUT4 translocation and glucose uptake in adipocytes, indicating that hyperactive kinase activity suppresses insulin-mediated glucose utilization through Rab8a and Rab10 phosphorylation (Kawakami et al. 2023; Imai et al. 2020).

Consistent with these findings, recent work has demonstrated that LRRK2 phosphorylates Rab8A in a glucose-dependent manner, facilitating its recruitment to the primary cilium (Dule et al. 2025). Our earlier study showed that the LRRK2-I1371V mutation drives kinase hyperactivity, resulting in Rab8A and Rab10 hyperphosphorylation (Potdar et al. 2024). Given that hyperphosphorylation of Rab8A and Rab10 has also been observed in LRRK2-I1371V astrocytes, the cell surface GLUT1 expression and glucose metabolism in these cells requires evaluation.

Organellar stress and dysfunction in PD has now been studied extensively, with the focus being centred around mitochondria, endoplasmic reticulum (ER) and lysosomes.

Mitochondrial abnormalities manifesting as loss of membrane potential and elevated reactive oxygen species (ROS) are well-recognized features of PD pathology (Mortiboys et al., 2010; Yue et al., 2015; Toyofuku et al., 2020; Wauters et al., 2020; Bonello et al., 2019; Korecka et al., 2019; Walter et al., 2019; Chen et al., 2022; Williamson et al., 2023). The intimate connection between mitochondrial function and cellular energy metabolism makes mitochondria particularly vulnerable to glucose deprivation and metabolic stress, conditions that can trigger mitochondrial permeability transition, cytochrome c release, and apoptotic cell death (Liu et al. 2007).

The ER serves as the primary site for protein folding, lipid synthesis, and calcium storage, making it exquisitely sensitive to cellular energy status and metabolic perturbations (Schwarz et al. 2015). ER stress, characterized by accumulation of misfolded proteins and activation of the unfolded protein response (UPR), has emerged as a significant pathogenic mechanism in PD, with evidence of ER stress markers in patient brain tissue and cellular models (Costa et al. 2020; Mou et al. 2020). Energy depletion compromises ER protein folding capacity through reduced ATP availability for chaperone function and impaired calcium homeostasis, leading to UPR activation and potential cell death if homeostasis cannot be restored (Malhotra et al. 2007).

Lysosomes function as the primary degradative organelles responsible for clearing damaged proteins, organelles, and other cellular debris through autophagy and endocytic pathways (Braulke et al. 2009; Saftig et al. 2009). Lysosomal dysfunction is intimately linked to PD pathogenesis, particularly given the association between α-synuclein accumulation and impaired protein clearance mechanisms (Dehay et al. 2013,51). The energy-intensive nature of lysosomal biogenesis, acidification, and proteolytic activity makes these organelles highly susceptible to ATP depletion and metabolic compromise, potentially creating a vicious cycle where energy deficits impair protein clearance, leading to further cellular dysfunction (Inpanathan et al. 2019).

Mounting evidence reveals extensive functional interconnections between mitochondria, ER, and lysosomes, collectively termed the "mitochondria-ER-lysosome axis," where dysfunction in one organelle can propagate to others through shared metabolic pathways, calcium signalling, and membrane contact sites (Todkar et al., 2017; Zhang et al., 2023). Mitochondria-ER contact sites facilitate calcium transfer, lipid synthesis, and metabolic coordination, while mitochondria-lysosome interactions regulate mitochondrial quality control through mitophagy (Lim et al. 2021; Wang et al. 2023). Similarly, ER-lysosome contacts support lipid transfer and stress responses, creating an integrated organellar network whose dysfunction can amplify cellular pathology (Lee et al. 2020).

In the context of energy deprivation, this organellar crosstalk can propagate metabolic stress throughout the cell, leading to coordinated organellar failure and cell death (Altman et al. 2012). Distinct LRRK2 variants have been associated with diverse organellar stresses, including impaired mitochondrial dynamics (Angeles et al., 2011; Sonninen et al., 2020), ER dysfunction affecting protein folding and cellular stress responses (Lee et al., 2019; Yao et al., 2023), and defective lysosomal degradation (Di Domenico et al., 2019; Lee et al., 2020). However, most studies have examined these effects in isolation, without considering how variant-specific disruptions may propagate across organellar networks to shape disease phenotypes.

Despite growing recognition of astrocytic contributions to LRRK2-associated PD, our understanding of how LRRK2 mutations, particularly the I1371V variant, affect astrocytic glucose metabolism and downstream organellar function remains limited. While previous studies have identified metabolic alterations in LRRK2 astrocytes, including changes in polyamine metabolism and mitochondrial respiration (Sonninen et al. 2020), the mechanistic links between glucose uptake deficits, energy failure, and multi-organellar dysfunction remain poorly characterized. Understanding these connections is crucial given that astrocytes provide essential metabolic support to neurons, and astrocytic energy failure may compromise neuronal survival and function in LRRK2-PD (Streubel-Gallasch et al. 2021).

Furthermore, the I1371V mutation, located in the Roc GTPase domain, may have distinct functional consequences compared to the more extensively studied G2019S kinase mutation, necessitating specific investigation of its effects on cellular metabolism and organellar health (Banerjee et al. 2023). This study therefore aims to understand how the LRRK2-I1371V mutation alters glucose uptake and metabolism in astrocytes and how these metabolic changes affect organellar function, particularly mitochondrial health, ER protein folding, and lysosomal degradative capacity. By linking LRRK2-I1371V-driven metabolic deficits to organellar dysfunction, we seek to uncover how astrocytic energy failure contributes to impaired neuronal support and vulnerability in PD.

### Methodology

**Reagents and tools table:**

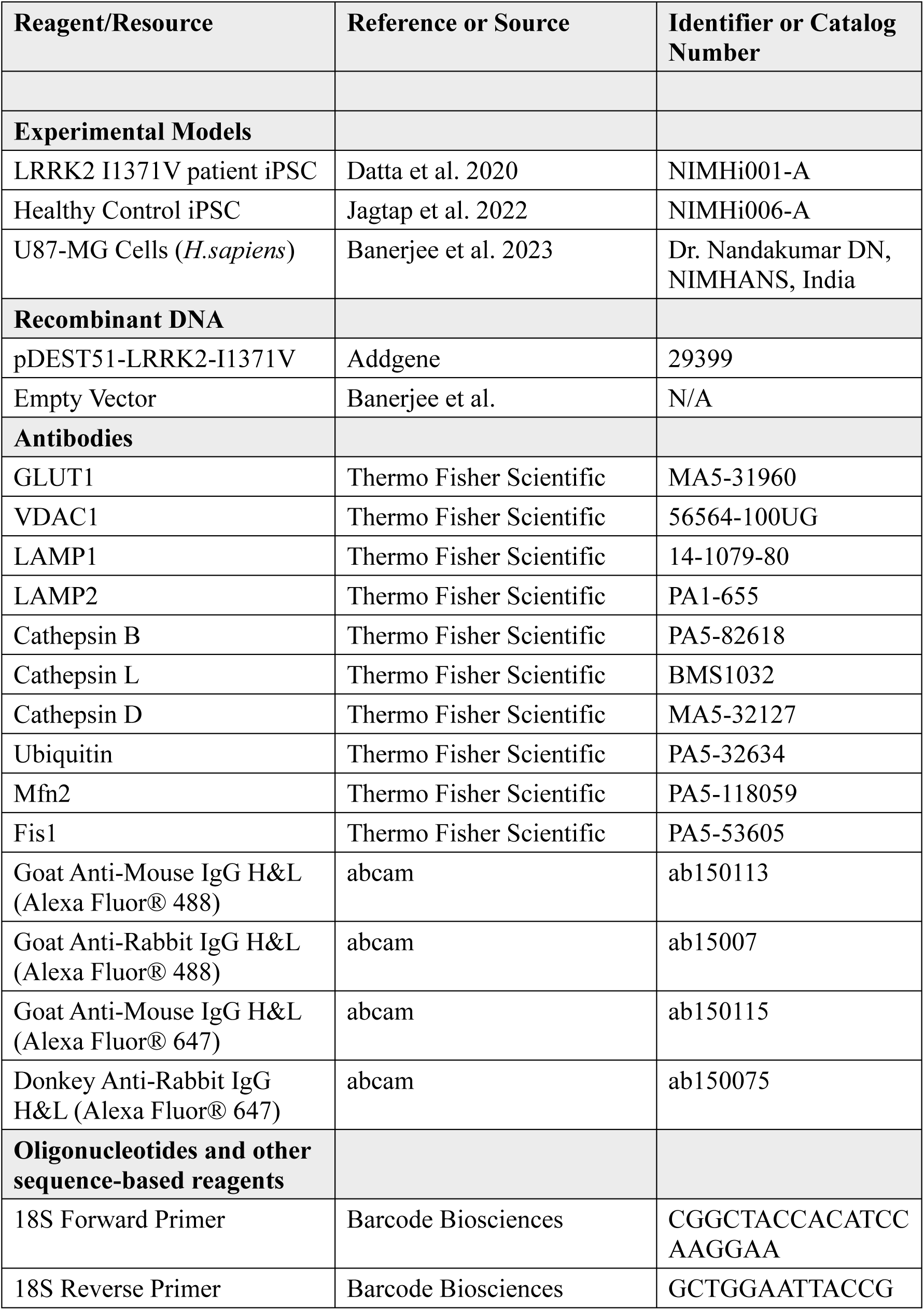
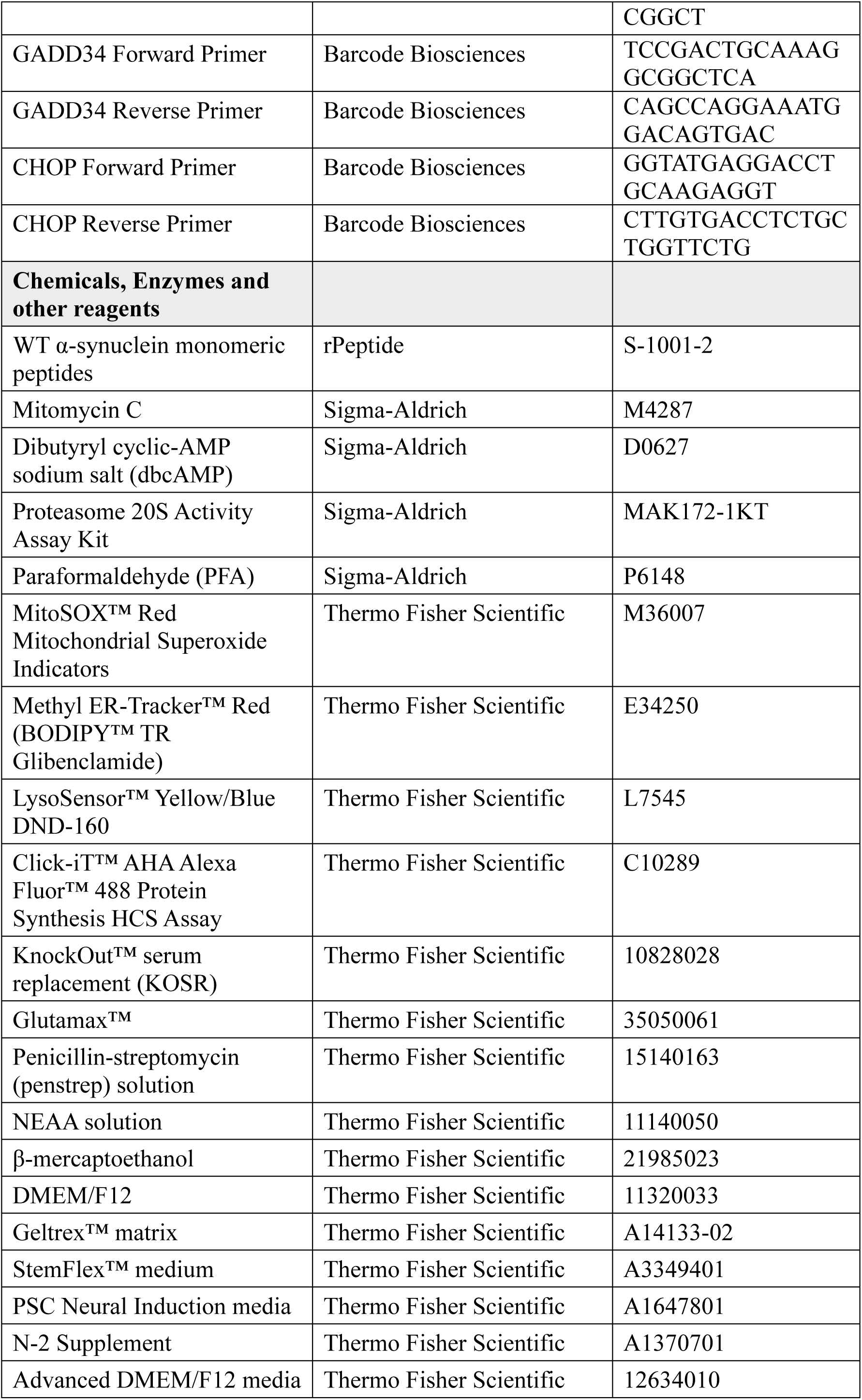
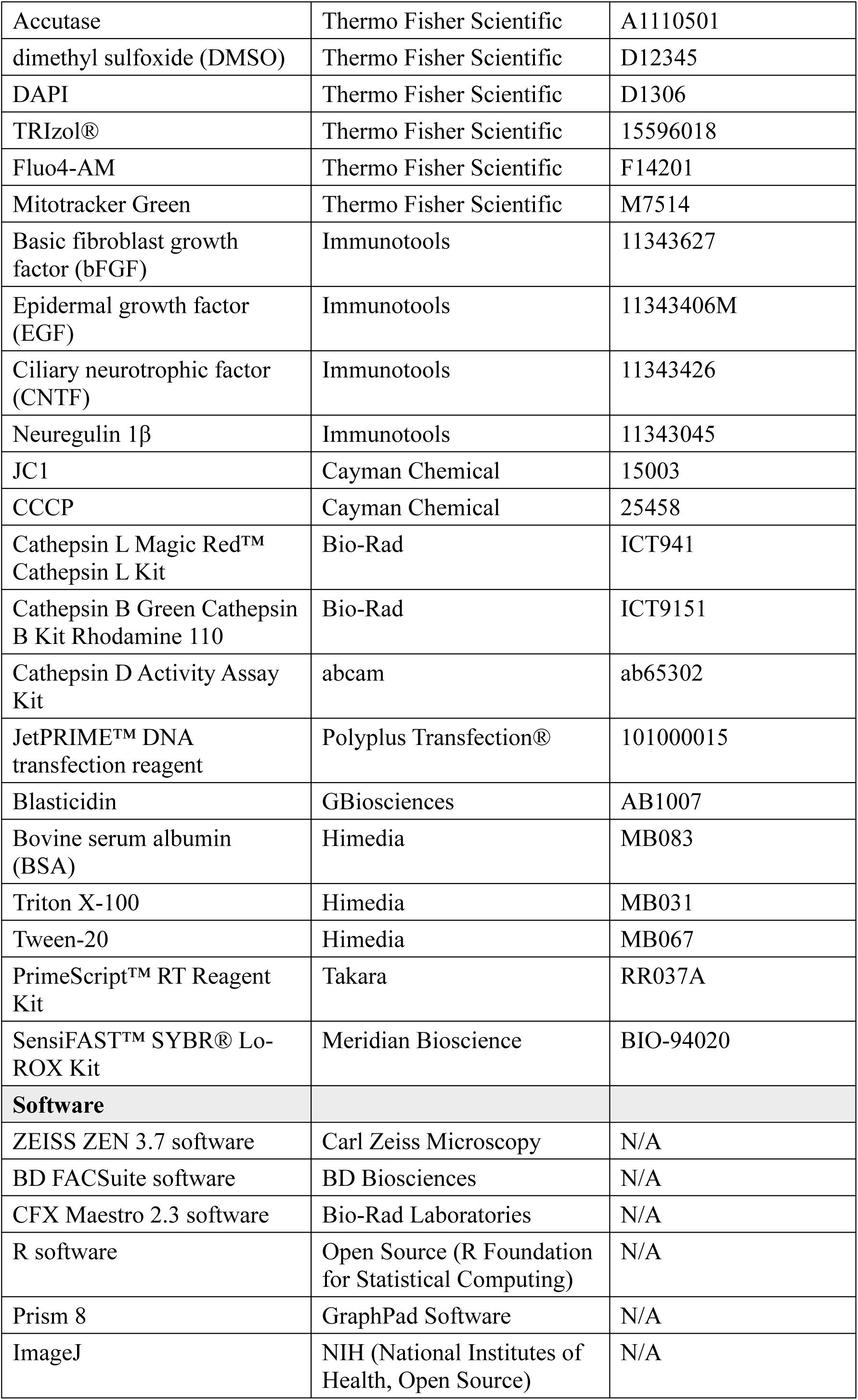
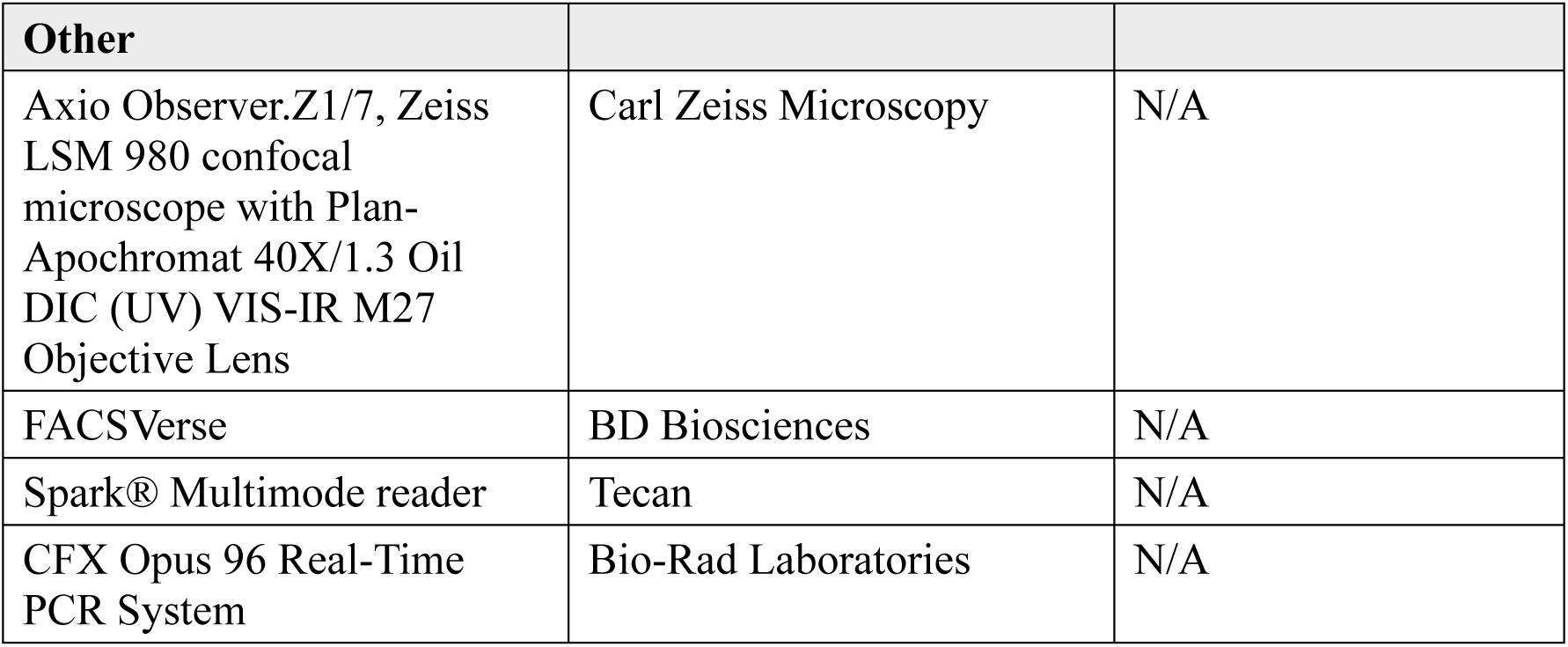

### Ethics Clearance

The generation, differentiation, and use of human induced pluripotent stem cells (hiPSCs) were approved by the Institutional Committee for Stem Cell Research (IC-SCR) under approval number SEC/05/030/BP. All animal experiments were performed in accordance with the guidelines of the Committee for the Purpose of Control and Supervision of Experiments on Animals (CPCSEA), Government of India, and were approved by the Institutional Animal Ethics Committee (IAEC) of the National Institute of Mental Health and Neuro Sciences (NIMHANS), reference number AEC/70/455/B.P. Animals were maintained under controlled environmental conditions with a 12 h light/dark cycle and ad libitum access to food and water.

### Cell Culture

Astrocytes were differentiated from hiPSCs using a two-step protocol optimized in our laboratory (Banerjee et al., 2023). Neural progenitor cells were first exposed to an astrocyte priming medium consisting of Neurobasal medium supplemented with N-2, GlutaMAX™, penicillin–streptomycin, EGF (10 ng/mL), and bFGF (10 ng/mL), thereby generating glial progenitor cells (GPCs) after four days. The cells were subsequently switched to a terminal differentiation medium composed of Neurobasal medium, N-2 supplement, GlutaMAX™, penicillin–streptomycin, CNTF (10 ng/mL), neuregulin-1β (10 ng/mL), and dbcAMP (0.1 µM) for three days, resulting in mature astrocytes. For comparison, the U87 MG glioblastoma cell line (kindly provided by Dr. Nandakumar DN, Department of Neurochemistry, NIMHANS) was cultured in DMEM/F12 supplemented with 10% fetal bovine serum (FBS), GlutaMAX™, and penicillin–streptomycin.

### Plasmid Constructs and Transfection

The pDEST51-LRRK2-I1371V (IV) plasmid containing a V5 tag was obtained from Mark Cookson via Addgene (plasmid #29399; RRID:Addgene_29399). An empty vector (EV) control was generated by HindIII digestion of the IV construct to remove the LRRK2 coding sequence (bp 291–6841). Transfections were performed as described previously (Banerjee et al., 2023). Briefly, U87 cells were transfected with EV or IV constructs using JetPRIME™ DNA transfection reagent. Stable transfectants were selected in medium containing 1.5 µg/mL blasticidin.

### Immunocytochemistry (ICC)

Adherent cells seeded on 12 mm coverslips at 90–95% confluency were fixed with 4% PFA. Permeabilization was performed using 1% Triton X-100, followed by blocking with 3% BSA. Primary antibodies (anti-GLUT1, anti-Ubiquitin, anti-VDAC1, anti-Mfn2, anti-Fis1, anti-LAMP1, anti-LAMP2, anti-Cathepsin B, anti-Cathepsin L, anti-Cathepsin D) were used at 1:100 dilution. Alexa Fluor® 488- and 647-tagged secondary antibodies were used at 1:200 dilution. Nuclei were stained with 300 nM DAPI. Washes were performed in PBS with 0.05% Tween-20. Confocal microscopy was carried out using a Zeiss LSM 980 with Airyscan super-resolution (Axio Observer.Z1/7) with a Plan Apochromat 40×/1.3 Oil DIC (UV) VIS-IR M27 objective. At least three biological replicates were analyzed with ≥10 fields imaged per condition. Image acquisition and analysis were performed using ZEISS ZEN 3.7 software, including generation of 2.5D views and calculation of Pearson’s correlation coefficients (n ≥ 5).

### Flow Cytometry

Astrocytes and U87 transfected cells were dissociated with StemPro™ Accutase™. Cells were fixed in 2% PFA for 45 min at room temperature, centrifuged at 10,000 rpm for 10 min (4 °C), and resuspended in DPBS with 0.01% sodium azide (1 × 10⁵ cells/reaction). For intracellular markers (GLUT1, Ubiquitin, VDAC1, Mfn2, Fis1, LAMP1, LAMP2, Cathepsin B, Cathepsin L, Cathepsin D), permeabilization was performed with 1% Triton X-100 for 15 min. Cells were blocked in 3% BSA for 45 min and incubated with primary antibodies (16 h, 4 °C), followed by fluorophore-tagged secondary antibodies (90 min, RT). For the surface expression of GLUT1, permeabilization step was omitted. Acquisition was performed on a BD FACSVerse flow cytometer with 10,000 events/sample.

### Lactate Production Assay

Lactate levels in the Conditioned Media was quantified using the L-Lactate Assay Kit (Elabscience) according to the manufacturer’s instructions. Absorbance was measured at 530 nm using an Infinite® 200 microplate reader (Tecan) to determine lactate release.

### Glucose Uptake Assay

Glucose uptake in HC and PD astrocytes was evaluated using the fluorescent D-glucose analogue 2-(N-(7-Nitrobenz-2-oxa-1,3-diazol-4-yl)Amino)-2-Deoxyglucose (2-NBDG; TCI). HC and PD astrocytes cultured on Geltrex coated plates, were washed with PBS to remove residual glucose before incubation. The cells were then exposed to 50 μM 2-NBDG in HBSS (without glucose) at 37 °C for 30 min. After incubation, they were washed twice with cold PBS, detached using Accutase, and centrifuged. The resulting cell pellet was resuspended in PBS, and fluorescence intensity was measured using a BD FACSLyric flow cytometer. Mean fluorescence intensity was compared between healthy control (HC) and Parkinson’s disease (PD) astrocytes to assess relative glucose uptake efficiency.

### Quantitative polymerase chain reaction (qPCR)

As described in our previous publications (Raj et al. 2022; Banerjee et al. 2023) qPCR was performed and analyzed using cDNA from treated and untreated HC and PD astrocytes. The primers used were 18S (Forward: 5’-CGGCTACCACATCCAAGGAA-3’, Reverse: 5’-GCTGGAATTACCGCGGCT-3’), GADD34 (Forward: 5’-TCCGACTGCAAAGGCGGCTCA-3’, Reverse: 5’- CAGCCAGGAAATGGACAGTGAC-3’) and CHOP (Forward: 5’- GGTATGAGGACCTGCAAGAGGT-3’, Reverse: 5’-CTTGTGACCTCTGCTGGTTCTG-3’). The qPCR was conducted using the CFX Opus 96 Real-Time PCR System (Bio Rad Laboratories), and data analysis was done with CFX Maestro 2.3 software. Results were represented as fold change in mRNA levels in PD and treated astrocytes, normalized to 18S mRNA and iNOS mRNA levels in Healthy Control (HC) astrocytes.

### Mitochondrial Membrane Potential

The mitochondrial membrane potential was determined in the control and peptide-treated cells (24 and 48 h) using the lipophilic cationic dye 5′,6,6′-tetrachloro-1,1′,3,3′-tetraethylbenzimidazolylcarbocyanine iodide (JC-1) as per the manufacturer’s instructions (Raj et al. 2023; Ganapathy et al. 2016). Dual analysis in the form of a bivariate plot was measured using FACS Verse (BD Biosciences) and analyzed using the software BD FACSuite. Additionally, red to green emission intensity ratio was plotted in the form of a graph.

### Mitotracker Green Staining

Mitochondrial mass was assessed using MitoTracker Green (MTG), which labels mitochondria in a membrane potential independent manner. Cells were incubated with 300 nM MTG and 300 nM DAPI, followed by confocal microscopy imaging.

### Endoplasmic Reticulum (ER)-Tracker™ Labeling and Imaging

Cells grown on 12-mm coverslips at ∼90–95% confluency were used for ER staining. All washes were performed with Hank’s balanced salt solution (HBSS; pH 7.4). Cells were incubated with 1 µM ER-Tracker™ Red following the manufacturer’s protocol and subsequently fixed with 4% paraformaldehyde. Imaging was performed on a Zeiss LSM 980 confocal microscope (Axio Observer.Z1/7) equipped with a Plan Apochromat 40×/1.3 Oil DIC (UV) VIS-IR M27 objective. For each condition, three independent samples were processed and a minimum of ten fields were acquired. Images were analyzed using ImageJ software (RRID:SCR_003070), and corrected total cell fluorescence (CTCF) was calculated from 50 regions of interest (ROIs) using the formula: CTCF = Integrated density − (Area of selected cell × Mean background fluorescence), as described previously (Raj et al., 2023; Sowmithra et al., 2020).

### Mitochondrial Superoxide Assay

Cells seeded on 12-mm coverslips at 90–95% confluency were stained with MitoSOX™ Red (Molecular Probes) according to the manufacturer’s instructions. After staining, cells were imaged on a Zeiss LSM 980 confocal microscope (Axio Observer.Z1/7) using the same optical configuration described above. Three biological replicates were analyzed, with at least ten fields captured per sample. Quantitative analysis was carried out in ImageJ (RRID:SCR_003070), and CTCF was calculated from 50 ROIs using the same formula employed for ER-Tracker fluorescence (Raj et al., 2023; Sowmithra et al., 2020).

### 20S Proteasome Activity Assay

Proteasome activity was assessed in 80,000 control and α-synuclein monomer-treated astrocytes seeded per well in a 96-well plate using the Proteasome 20S Activity Assay Kit, following the manufacturer’s instructions. Fluorescence intensity was measured at an excitation wavelength of 490 nm and an emission wavelength of 525 nm using an Infinite® 200 microplate reader (Tecan).

### Intracellular Cathepsin Activity Assays

Intracellular cathepsin activities were measured using the Magic Red™ Cathepsin L Detection Kit, the Green Cathepsin B Detection Kit, and the Cathepsin D Activity Assay Kit, following the manufacturers’ instructions. Cathepsin L and D activities were assessed using a fluorometric format on an Infinite® 200 microplate reader (Tecan) with excitation/emission wavelengths of 592/628 nm for Cathepsin L and 328/460 nm for Cathepsin D. Cathepsin B activity was measured in live cells using flow cytometry with an excitation/emission of 488/525 nm. All measurements were performed five times, and data were used to quantify enzyme activity in HC and PD astrocytes.

### Lysosomal pH Measurement

Lysosomal pH was determined using LysoSensor™ Yellow/Blue DND-160 (Thermo Fisher Scientific). Cells grown on coverslips at 90–95% confluency were stained with 1 µM probe in prewarmed growth medium for 5 min at 37 °C, as per manufacturer’s instructions. After incubation, the dye was replaced with fresh medium, and coverslips were mounted directly for imaging. Fluorescence was acquired on a Zeiss LSM 980 confocal microscope using the Plan Apochromat 40×/1.3 Oil DIC objective. For each condition, five independent samples were processed with ≥10 fields per replicate. Quantitative analysis of fluorescence was performed in ImageJ using CTCF from 50 ROIs as described earlier (Raj et al., 2023; Sowmithra et al., 2020).

### Transmission Electron Microscopy (TEM)

Cells from at least two 60-mm dishes (∼95% confluency) were processed for TEM as described previously (Raj et al. 2022; 2023). Cells were primarily fixed in 3% buffered glutaraldehyde for 1 hour, post fixed 1% osmium tetroxide for 1 hour. Sodium cacodylate buffer (0.1M, pH-7.2-7.4) was used for washes between fixation steps. Samples were then dehydrated through a graded ethanol series, cleared in propylene oxide, and embedded in Araldite CY212 resin, which was polymerized at 60 °C for 48 h. Ultrathin sections were cut using a Leica EM UC7 ultramicrotome (Leica Mikrosysteme, Austria) and contrasted with saturated methnolic uranyl acetate and lead citrate. Imaging was performed on a JEM-1400 Plus TEM (JEOL, Japan) operated at 80 kV, with images captured using a Gatan SC1000B camera (Santhoshkumar et al. 2021).

### Statistics

All results are expressed as mean ± SD. Statistical comparisons were performed using two-sample t-Test or one-way ANOVA, as appropriate, followed by Bonferroni post hoc analysis, using R software (R Foundation; R Project for Statistical Computing). A p-value less than 0.05 was considered significant. Graphs were created using GraphPad Prism 8 (GraphPad Software). Unless otherwise stated graphed data are presented as means ± SD. Single symbol: P<0.05; double-symbol: P<0.01 triple-symbol: P<0.001.

## Results

### Disrupted glucose handling and reduced lactate release in PD astrocytes

Astrocytes are the primary regulators of neuronal energy supply. They import glucose predominantly through the transporter GLUT1 at their end foot processes convert it via glycolysis into lactate, and shuttle lactate to neurons through MCT4 to sustain oxidative metabolism (Gonçalves et al. 2019). Disruption of this astrocyte neuron lactate shuttle has been increasingly linked to neurodegeneration in PD (Singh et al. 2025).

To investigate whether glucose metabolism is impaired in I1371V astrocytes, we differentiated iPSCs from a healthy control (HC) and a PD patient carrying the I1371V mutation into astrocytes using our previously published protocol (Banerjee et al., 2023). Immunofluorescence analysis showed significantly reduced GLUT1 expression in PD astrocytes compared to controls (Fig. 1A–B, p<0.001). Flow cytometric analysis corroborated this finding, demonstrating a markedly lower percentage of GLUT1-immunopositive cells in the PD astrocyte population relative to HC astrocytes (Fig. 1C, Supplementary Fig. 1A-B p<0.001). To validate these observations in an independent model system, we transfected U87 cells with either LRRK2 I1371V (IV) or empty vector (EV) controls. Consistent with our iPSC-derived astrocyte data, IV-transfected U87 cells exhibited a significantly lesser GLUT1-immunopositive population compared to EV-transfected controls (Fig. 1D, Supplementary Fig. 1C-D, p<0.001). Analysis of cell surface GLUT1 expression revealed significantly diminished plasma membrane localization in PD astrocytes compared to HC controls (Fig. 1E, Supplementary Fig. 1E-F p<0.001), a finding that was reproducibly observed in the U87 transfection model (Fig. 1F, Supplementary Fig. 1G-H; p<0.001). Since decreased GLUT1 expression is directly linked to compromised glucose transport, we next evaluated 2-NBDG uptake, a fluorescent glucose analogue, using flow cytometry (Fig. 1G). PD astrocytes exhibited significantly lower 2-NBDG uptake relative to HC astrocytes (Fig. 1H, p<0.001). This reduction in glucose uptake was further underlined by lactate measurements in the conditioned medium, which showed that PD astrocytes released substantially less lactate than their HC counterparts (Fig. 1I, p < 0.001). Collectively, these findings along with our earlier report of lesser ATP in PD LRRK2 I1371V astrocytes (Banerjee et al., 2023) demonstrates impaired glucose uptake can be one of the primary reasons for lesser ATP levels.

**Figure 1.**
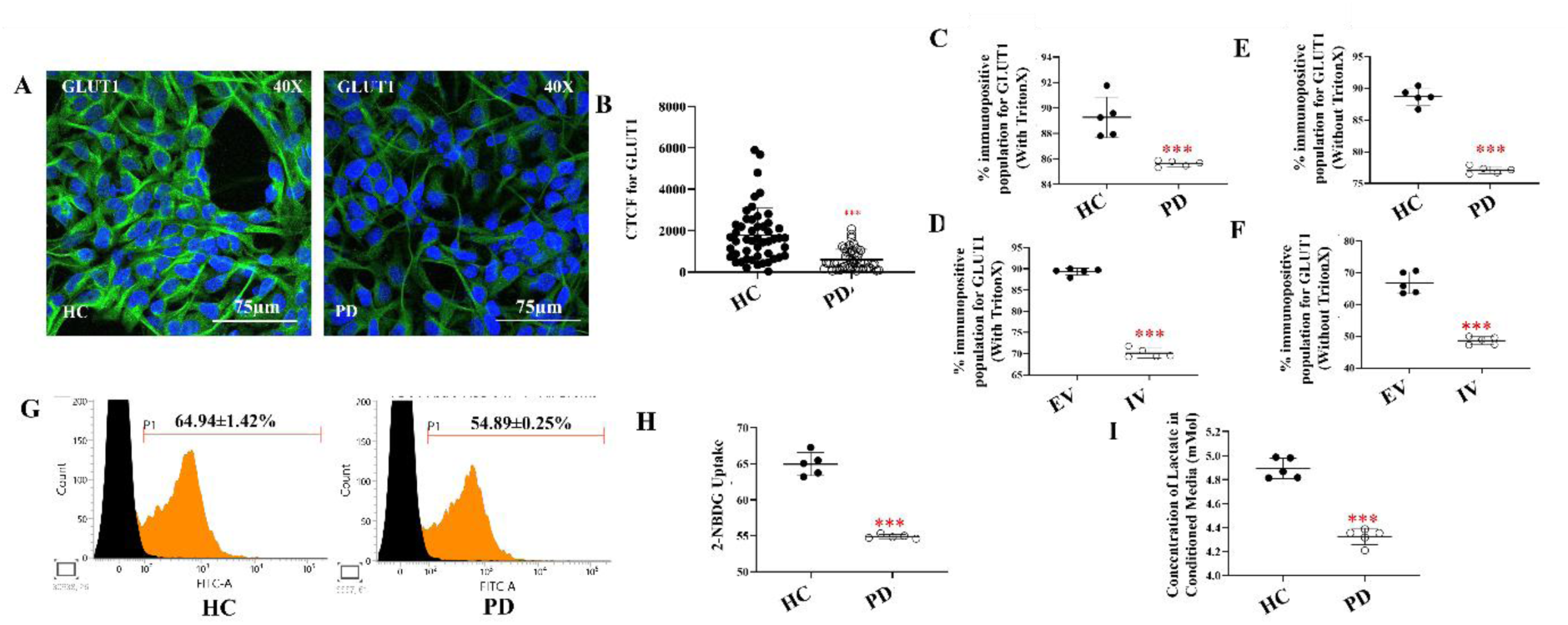
Impaired Glucose Handling in LRRK2 I1371V Astrocytes. (A) ICC images of HC and PD astrocytes stained for GLUT1, with nuclei counterstained with DAPI. (B) Scatter plot of the CTCF values of the intensity of GLUT1 [p<0.001; HC vs PD] (Data are mean ± SD, Two Sample t-Test; n=50). (C-D) Graphical representation of the flow cytometry quantified positive population for total GLUT1 in HC and PD astrocytes (C) and EV and IV transfected U87 cells (D). (Data are mean ± SD, Two Sample t-Test; n=5). [p<0.001; HC vs PD; EV vs IV] (E-F) Graphical representation of the flow cytometry quantified positive population for Membrane localized GLUT1 in HC and PD astrocytes (E) and EV and IV transfected U87 cells (F). (Data are mean ± SD, Two Sample t-Test; n=5). [p<0.001; HC vs PD; EV vs IV](G) Representative flow cytometry histograms of 2-NBDG uptake in HC and PD astrocytes. (H) Quantification of mean fluorescence intensity (MFI) from 2-NBDG flow cytometric assay. [p<0.001; HC vs PD] (Data are mean ± SD, Two Sample t-Test; n=5). (I) Release of Lactate into the Conditioned Media from HC and PD Astrocytes. [p<0.001; HC vs PD] (Data are mean ± SD, Two Sample t-Test; n=5).

### Loss of Mitochondrial Membrane Potential, Elevated ROS, and Mitophagy Signals in PD Astrocytes

The other contributory factor for the lesser ATP levels (Jouaville et al. 1999) can be due to mitochondrial impairment, as ATP, the final product of glucose metabolism, is primarily generated within mitochondria. We assessed mitochondrial membrane potential using the ratiometric JC-1 dye, which measures the red-to-green fluorescence ratio. Flow cytometry analysis demonstrated that healthy control (HC) astrocytes exhibited a significantly higher red/green fluorescence ratio compared to PD astrocytes, indicating more intact mitochondrial membrane potential (Fig. 2A–B). Representative event scatter plots are shown in Fig. 2A, with corresponding quantitative analysis presented in Fig. 2B. Treatment with CCCP caused a marked decrease in the red/green ratio in HC astrocytes, confirming assay validity.

**Figure 2.**
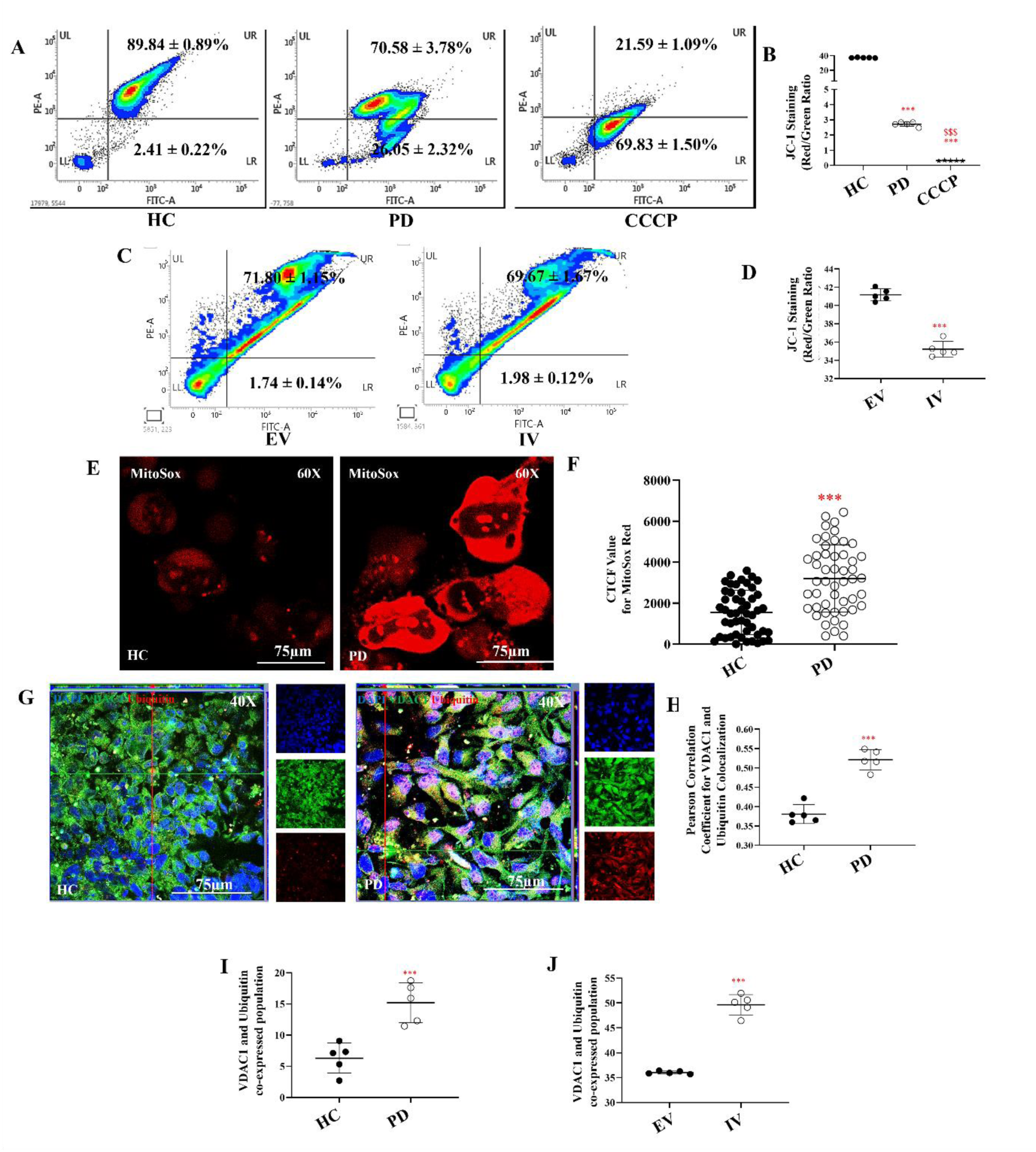
Mitochondrial Impairment in LRRK2 I1371V Astrocytes. (A) Representative JC-1 flow cytometry histograms illustrating red (aggregate) to green (monomer) fluorescence in HC, PD and CCCP-treated positive control astrocytes. (B) Quantification of JC-1 aggregate/monomer ratios in HC, PD and CCCP-treated positive control astrocytes [p<0.001; HC vs PD; CCCP; PD vs CCCP] (Data are mean ± SD, One Way ANOVA; n=5). (C) Representative JC-1 flow cytometry histograms illustrating red (aggregate) to green (monomer) fluorescence in EV and IV transfected U87 cells (D) Quantification of JC-1 aggregate/monomer ratios EV and IV transfected U87 cells. [p<0.001; EV vs IV] (Data are mean ± SD, Two Sample t-Test; n=5). (E) Representative confocal images of cells stained with MitoSOX Red to detect mitochondrial ROS levels in HC and PD astrocytes. (F) ) Scatter plot of the CTCF values of the intensity of MitoSOX Red [p<0.001; HC vs PD] (Data are mean ± SD, Two Sample t-Test; n=50). (G) ICC images of VDAC1 (Alexa Fluor^®^ 488; green) and Ubiquitin (Alexa Fluor^®^ 647; red) coimmunostained HC and PD astrocytes; nucleus counterstained with DAPI. (H) Pearson’s coefficient of the colocalization of VDAC1 with Ubiquitin. [p<0.001; HC vs PD] (Data are mean ± SD, Two Sample t-Test; n=5) (I-J) Graphical representation of percentage of co-positive population of VDAC1 with Ubiquitin in HC and PD astrocytes (I) and EV and IV transfected U87 cells (J). [p<0.001; HC vs PD; EV vs IV] (Data are mean ± SD, Two Sample t-Test; n=5).

We obtained similar results in U87 cells transfected with LRRK2 I1371V versus empty vector (EV) controls. FACS analysis revealed that cells expressing the I1371V variant displayed significantly reduced mitochondrial membrane potential compared to EV controls (Fig. 2C–D). To ensure reproducibility, we repeated the JC-1 assay using an independent iPSC-derived astrocyte line from a second healthy control (HC02), which yielded comparable results (Supplementary Fig. 2 A-B).

We next investigated whether the mutation also affected mitochondrial oxidative stress. Using MitoSOX Red, we observed markedly elevated fluorescence in PD astrocytes compared to HC cells, indicating increased mitochondrial ROS levels (Fig. 2E). Corrected total cell fluorescence (CTCF) analysis confirmed a significant increase in mitochondrial ROS in PD astrocytes (p < 0.001, Fig. 2G).

Given that mitochondria in PD astrocytes experience significant stress, we examined whether these compromised organelles were being targeted for clearance through mitophagy—a critical protective mechanism for maintaining cellular homeostasis. To confirm that mitochondria in PD astrocytes were indeed stressed and primed for clearance, we assessed ubiquitin recruitment to these organelles. Co-localization analysis of ubiquitin (Ubq) with VDAC1, an outer mitochondrial membrane marker, revealed increased yellow puncta in PD astrocytes (Fig. 2G), indicating enhanced mitochondrial ubiquitination. Pearson’s coefficient analysis substantiated this observation, showing significantly higher Ubq–VDAC1 co-localization in PD cells (Fig. 2H; p < 0.001). Flow cytometric analysis corroborated these findings, with PD astrocytes displaying a significantly higher proportion of co-labelled cells compared to HC (Fig. 2I, Supplementary Fig. 2C-D; p < 0.001). LRRK2 I1371V-transfected cells recapitulated these results, exhibiting markedly increased Ubq–VDAC1 co-labelling compared to EV controls (Fig. 2J, Supplementary Fig. 2E-F; p < 0.001).

We next asked whether mitochondrial stress in PD astrocytes was linked to changes in mitochondrial dynamics. To address this, we examined key proteins involved in mitochondrial fission and fusion. Immunofluorescence staining of the fission protein FIS1 (Fig. 3A) and the fusion protein MFN2 (Fig. 3B) showed similar patterns in HC and PD astrocytes. Quantification using CTCF revealed no significant differences in protein levels (Fig. 3C–D; p > 0.05). Flow cytometry confirmed these findings, with no detectable differences between groups (Fig. 3E–F; Supplementary Fig. 2M–P).

**Figure 3:**
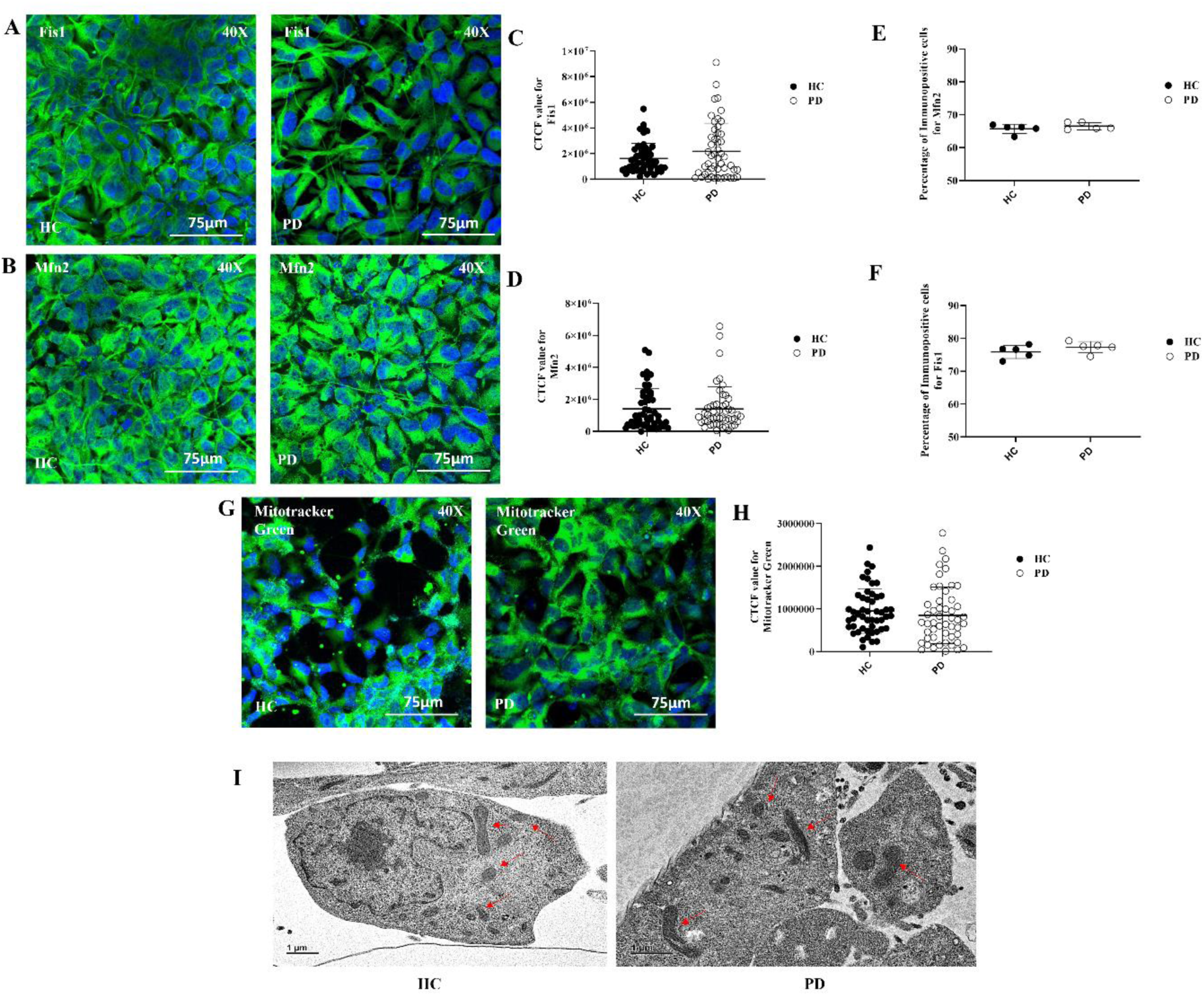
Mitochondrial fission–fusion dynamics and ultrastructure are preserved in LRRK2 I1371V astrocytes. ICC images of HC and PD astrocytes stained for Fis1 (A) and Mfn2 (B), with nuclei counterstained with DAPI. (C-D) Scatter plot of the CTCF values of the intensity Fis1 (E) and Mfn2 (F) [p>0.05; HC vs PD] (Data are mean ± SD, Two Sample t-Test; n=50). (G-H) Graphical representation of the flow cytometry quantified positive population for Fis1 (F) and Mfn2 (G) in HC and PD astrocytes [p>0.05; HC vs PD] (Data are mean ± SD, Two Sample t-Test; n=5). (I) Transmission electron microscopy (TEM) images depict mitochondrial ultrastructure following 24-hour treatment with wild-type α-synuclein. Red arrows indicate mitochondria.

With mitochondrial dynamics unchanged, we next examined mitochondrial mass. Cells were stained with MitoTracker Green (Fig. 3G). Quantitative analysis showed no difference in fluorescence intensity between HC and PD astrocytes (Fig. 3H; p > 0.05). This indicated that mitochondrial mass was preserved. Ultrastructural analysis was performed using TEM. Mitochondria in HC and PD astrocytes appeared similar. No significant changes in morphology or cristae organization were observed (Fig. 3I).

We next asked whether mitochondrial stress in PD astrocytes was linked to changes in mitochondrial dynamics. To address this, we examined key proteins involved in mitochondrial fission and fusion. Immunofluorescence staining of the fission protein FIS1 (Fig. 3A) and the fusion protein MFN2 (Fig. 3B) showed similar patterns in HC and PD astrocytes. Quantification using CTCF revealed no significant differences in protein levels (Fig. 3C–D; p > 0.05). Flow cytometry confirmed these findings, with no detectable differences between groups (Fig. 3E–F; Supplementary Fig. 2G–J).

With mitochondrial dynamics unchanged, we next examined mitochondrial mass. Cells were stained with MitoTracker Green (Fig. 3G). Quantitative analysis showed no difference in fluorescence intensity between HC and PD astrocytes (Fig. 3H; p > 0.05). This indicated that mitochondrial mass was preserved. Ultrastructural analysis was performed using TEM. Mitochondria in HC and PD astrocytes appeared similar. No significant changes in morphology or cristae organization were observed (Fig. 3I).

### Proteasomal Dysfunction and Lysosomal Deficits in I1371V Astrocytes

Given the compromised mitochondrial function and reduced ATP synthesis, we investigated how this metabolic vulnerability affects protein degradation—a highly energy-dependent process. We examined the proteasome, which plays an essential role in degrading ubiquitinated mitochondria during mitophagy. Assessment of 20S proteasomal activity revealed significant reduction in PD astrocytes compared to HC controls (Fig. 4A; p < 0.01). These findings indicate that although dysfunctional mitochondria in PD astrocytes are appropriately tagged for clearance, impaired proteasomal activity prevents their efficient removal, thereby contributing to the persistence of stressed organelles and exacerbating cellular dysfunction.

**Figure 4.**
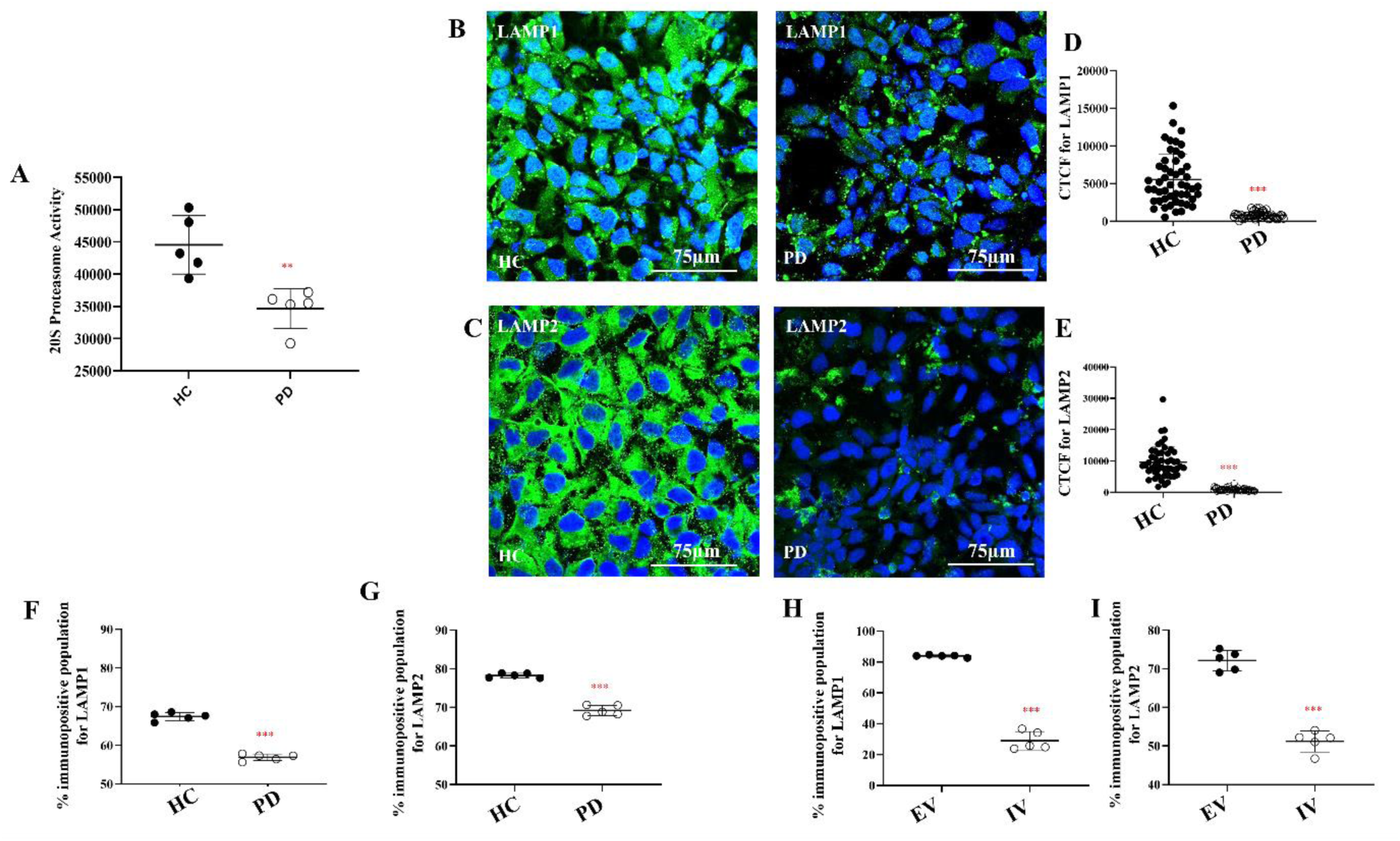
Decreased Proteasomal Activity and Reduction of the Number of Lysosomes in LRRK2 I1371V Astrocytes. (A) Quantification of 20S proteasome activity in HC and PD Astrocytes [p<0.001; HC vs PD] (Data are mean ± SD, Two Sample t-Test; n=5). (B-C) ICC images of HC and PD astrocytes stained for LAMP1 (B) and LAMP2 (C), with nuclei counterstained with DAPI. (D-E) Scatter plot of the CTCF values of the intensity LAMP1 (D) and LAMP2 (E) [p<0.001; HC vs PD] (Data are mean ± SD, Two Sample t-Test; n=50). (F-G) Graphical representation of the flow cytometry quantified positive population for LAMP1 (F) and LAMP2 (G) in HC and PD astrocytes [p<0.001; HC vs PD] (Data are mean ± SD, Two Sample t-Test; n=5). (H-I) Graphical representation of the flow cytometry quantified positive population for LAMP1 (H) and LAMP2 (I) in EV and IV transfected U87 cells [p<0.001; EV vs IV] (Data are mean ± SD, Two Sample t-Test; n=5).

Following the identification of proteasomal impairment, we examined the lysosomal degradation pathway, which represents the final step in autophagic clearance. Immunocytochemistry revealed reduced expression of lysosomal markers LAMP1 and LAMP2 in PD astrocytes compared to HC controls (Fig. 4B-C). Quantitative corrected total cell fluorescence (CTCF) analysis confirmed significant decreases in both markers (Fig. 4D–E; p < 0.001). Flow cytometry validated these findings, demonstrating significantly lower LAMP1 and LAMP2 expression in PD astrocytes (Fig. 4F,G, Supplementary Fig. 3A-D; p < 0.001). Importantly, no significant differences were observed between two independent healthy control astrocyte lines (HC01 and HC02) (Supplementary Fig. 3 E-H), confirming the mutation-specific nature of these changes. Supporting this conclusion, I1371V-transfected U87 cells similarly exhibited reduced LAMP1 and LAMP2 expression compared to empty vector controls (Fig. 4H–I, Supplementary Fig. 3I-L; p < 0.001).

To further investigate lysosomal functionality, we examined the expression of key lysosomal proteases: Cathepsins B, L, and D. ICC analysis revealed no observable differences in fluorescence intensity for Cathepsins B and L between PD and HC astrocytes (Fig. 5A–B). In contrast, Cathepsin D fluorescence was significantly reduced in PD astrocytes (Fig. 5C). CTCF quantification confirmed these observations, showing no significant differences for Cathepsins B and L between groups (Fig. 5D–E), while Cathepsin D levels were significantly lower in PD astrocytes (Fig. 5F; p < 0.001). Flow cytometry corroborated these findings, with Cathepsins B and L expression remaining comparable between HC and PD groups (Fig. 5G–H, Supplementary Fig. 4A-D), whereas Cathepsin D expression was significantly decreased in PD astrocytes (Fig. 5I, Supplementary Fig. 4E-F; p < 0.001). Parallel analysis of EV and IV cells yielded identical results, highlighting reduced Cathepsin D expression relative to EV controls (Fig. 5J–L, Supplementary Fig. 4G-L; p < 0.001).

**Figure 5.**
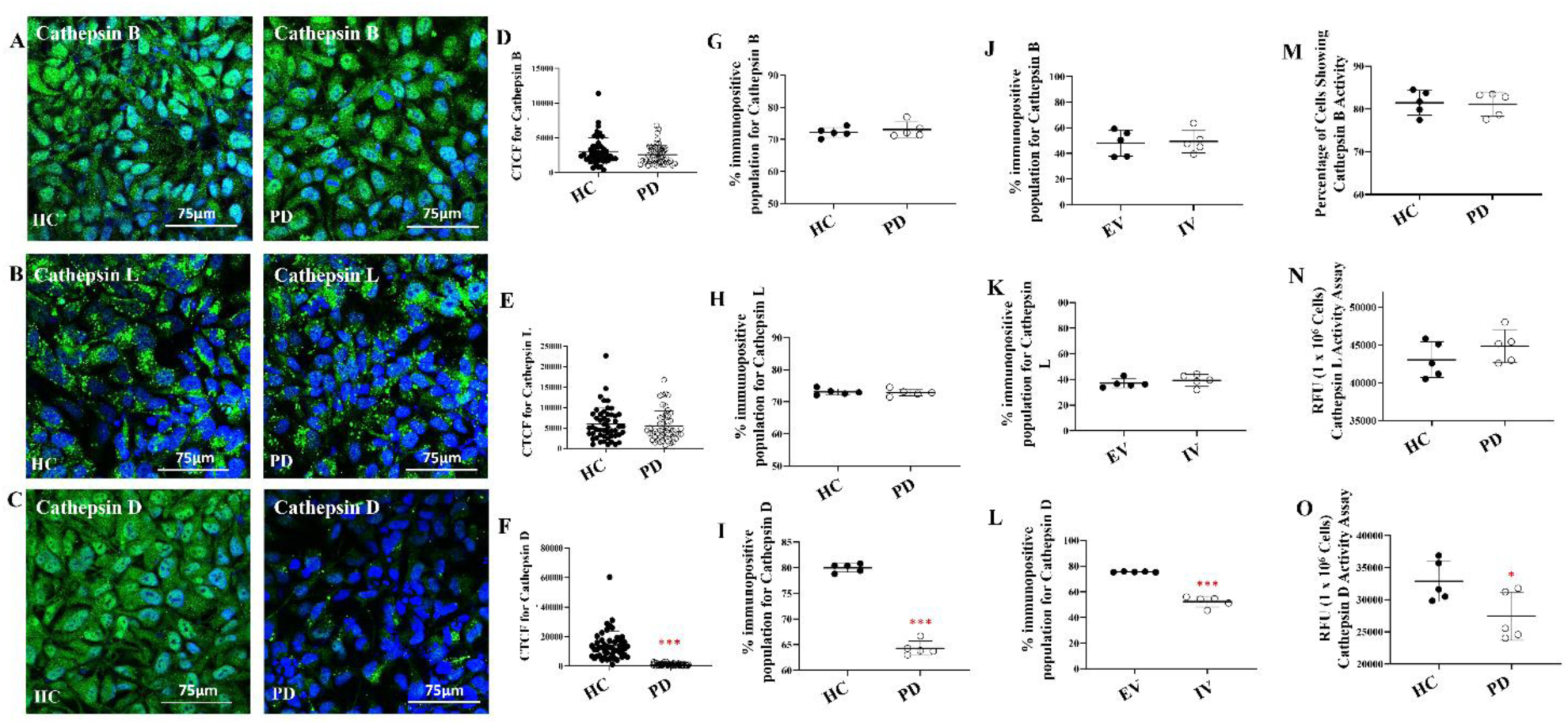
Selective Reduction Cathepsin D Levels and Activity in LRRK2 I1371V Astrocytes. (A-C) ICC images of HC and PD astrocytes stained for Cathepsin B (A), Cathepsin L (B) and Cathepsin D (C) with nuclei counterstained with DAPI. (D-F) Scatter plot of the CTCF values of the intensity Cathepsin B (D), Cathepsin L (E) and Cathepsin D (F) [p<0.001; HC vs PD] (Data are mean ± SD, Two Sample t-Test; n=50). (G-I) Graphical representation of the flow cytometry quantified positive population for Cathepsin B (G), Cathepsin L (H) and Cathepsin D (I) in HC and PD astrocytes (G) [p<0.001; HC vs PD] (Data are mean ± SD, Two Sample t-Test; n=5). (J-L) Graphical representation of the flow cytometry quantified positive population for Cathepsin B (J), Cathepsin L (K) and Cathepsin D (L) in EV and IV transfected U87 cells (I) [p<0.001; EV vs IV] (Data are mean ± SD, Two Sample t-Test; n=5). (M) Cathepsin B Activity in HC and PD Astrocytes. (Data are mean ± SD, Two Sample t-Test; n=5). (N) Cathepsin L Activity in HC and PD Astrocytes. (Data are mean ± SD, Two Sample t-Test; n=5). (O) Cathepsin D Activity in HC and PD Astrocytes. (Data are mean ± SD, Two Sample t-Test; n=5).

To determine whether these expression changes translated to functional differences, we assessed enzymatic activities of these cathepsins. Flow cytometry-based activity assays for Cathepsin B and spectrofluorometric assays for Cathepsin L revealed comparable enzymatic activities between healthy control and PD astrocytes, with no statistically significant differences observed (Fig. 5M–N; Supplementary Fig. 4M-N). In stark contrast, functional assessment of Cathepsin D enzymatic activity revealed a significant reduction in PD astrocytes compared to healthy controls (Fig. 5O; p < 0.001), demonstrating that both Cathepsin D expression and its proteolytic function are compromised in these cells.

To gain additional insight into lysosomal function, we assessed lysosomal pH using LysoSensor-based immunofluorescence imaging. HC astrocytes displayed robust LysoSensor fluorescence, while PD astrocytes exhibited significantly reduced signal intensity (Fig. 6A–B; p < 0.001), consistent with impaired acidification. Since acidic pH is essential for lysosomal protease function, this finding suggested compromised degradative capacity. To test this hypothesis, we exposed astrocytes to wild-type monomeric α-synuclein for 24 hours and examined their ultrastructure using transmission electron microscopy (TEM). HC astrocytes displayed lysosomes containing clearly degraded material, whereas PD astrocytes showed numerous lysosomes packed with undegraded vesicular content (Fig. 6C).

**Figure 6.**
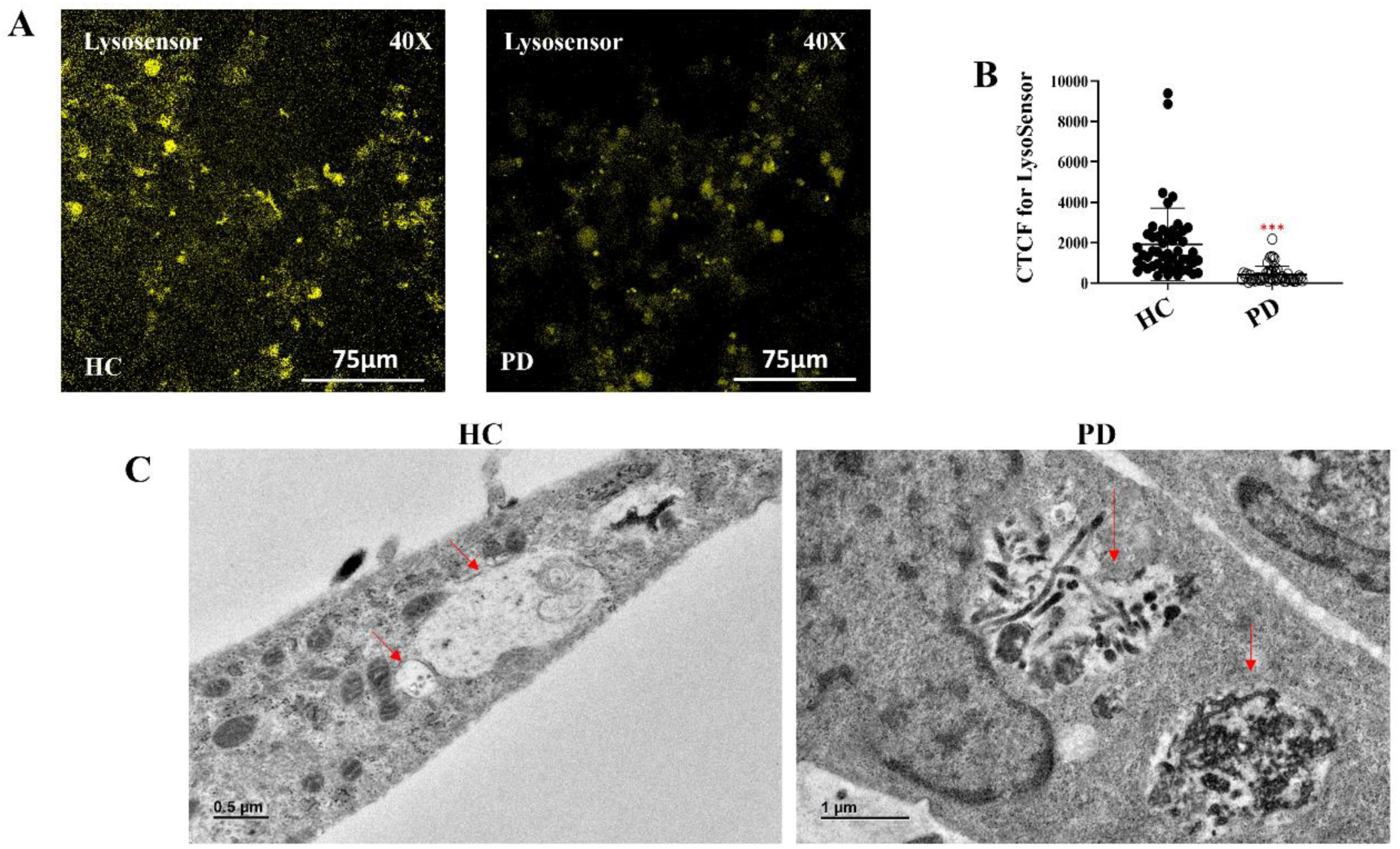
Altered Lysosomal pH and Dense Lysosomal Accumulation upon α-Synuclein Treatment in in LRRK2 I1371V Astrocytes. (A) Representative confocal images of LysoSensor staining showing lysosomal pH in healthy control (HC) and LRRK2 I1371V Parkinson’s disease (PD) astrocytes. (B) Scatter plot of the CTCF values of the intensity of Lysosensor [p<0.001; HC vs PD] (Data are mean ± SD, Two Sample t-Test; n=50). (C) Transmission electron microscopy (TEM) images depict lysosomal ultrastructure following 24-hour treatment with wild-type α-synuclein. Red arrows indicate lysosomes. PD astrocytes display densely packed, electron-dense lysosomes, reflecting an accumulation of undigested material, whereas HC astrocytes exhibit normal lysosomal morphology.

These findings demonstrate that the LRRK2 I1371V mutation compromises lysosomal integrity in PD astrocytes. The combination of reduced lysosome number, impaired acidification, and diminished Cathepsin D activity collectively hinders autophagy. Consequently, damaged mitochondria and undegraded cargo accumulate, perpetuating mitochondrial stress and driving astrocytic dysfunction.

### ER Stress and Impaired Protein Synthesis in LRRK2 I1371V Astrocytes

Energy deficits arising from mitochondrial dysfunction and impaired lysosomal clearance are expected to hamper energy-intensive cellular processes, including protein synthesis. Building on the evidence of mitochondrial depolarization and oxidative stress, the LRRK2 I1371V mutation was hypothesized to perturb ER function, given its central role in protein folding, calcium homeostasis, and organelle cross talk. To assess ER integrity, astrocytes were stained with ER Tracker dye. PD astrocytes exhibited significantly reduced fluorescence intensity compared to healthy controls (HC), suggesting ER disruption and elevated stress (Fig. 7A). Quantification using CTCF confirmed a significant reduction in ER signal in PD cells (Fig. 7B; p<0.001). ER stress was further examined at the transcriptional level by measuring the expression of key unfolded protein response (UPR) genes. Both GADD34 and CHOP genes were significantly upregulated in PD astrocytes in comparison to HC, leading to activation of the ER stress response (Fig. 7C–D; p<0.001). At the protein level, phosphorylated PERK (p-PERK) and CHOP levels were assessed using flow cytometry. Both markers were significantly higher in PD astrocytes compared to HC (Fig. 7E–F, Supplementary Fig. 5A-D; p<0.001). To ensure that these differences were mutation specific and not due to variability among control lines, a second healthy iPSC derived astrocyte line (HC02) was analyzed, which showed no significant difference in expression compared to the original HC line (Supplementary Fig. 5E–H). Similar upregulation of p-PERK and CHOP was observed in U87 cells transfected with EV and IV, with IV cells showing significantly higher levels of both markers (Fig. 7G–H, Supplementary Fig. 5I-L; p<0.001). This confirmed that the LRRK2 I1371V mutation directly contributes to ER stress activation. As chronic ER stress is known to impair protein synthesis, nascent protein production was evaluated using the Click-iT assay. Consistent with elevated stress levels, PD astrocytes exhibited a significant reduction in protein synthesis compared to HC, as quantified by CTCF (Fig. 7I–J; p<0.001), further supporting the presence of ER dysfunction in LRRK2 mutant astrocytes.

**Figure 7.**
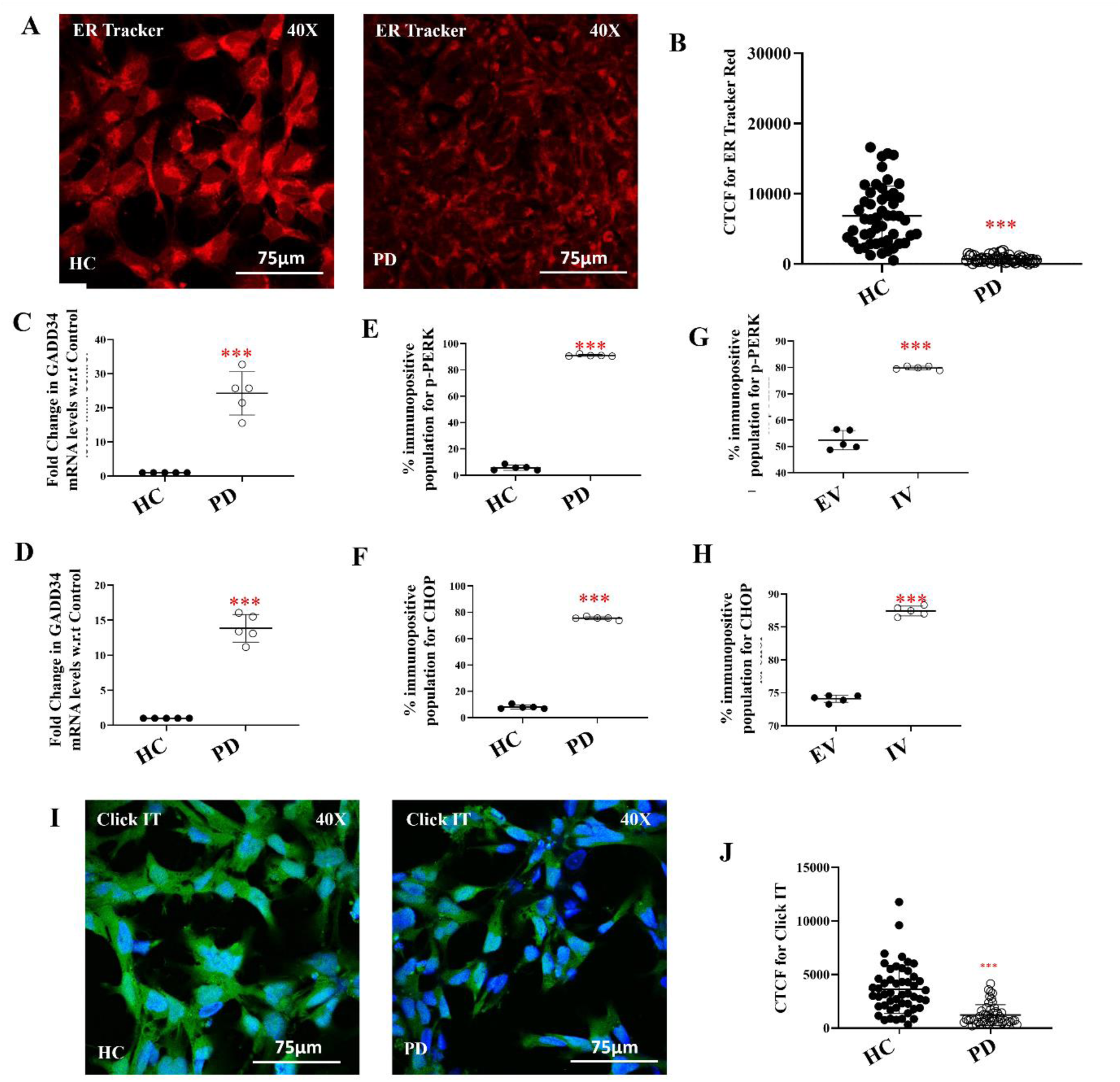
Altered Lysosomal pH and Dense Lysosomal Accumulation upon α-Synuclein Treatment in LRRK2 I1371V Astrocytes. (A) Representative confocal images showing ER morphology using methyl ER-Tracker™ Red (BODIPY™ TR Glibenclamide) staining in healthy control (HC) and LRRK2 I1371V Parkinson’s disease (PD) astrocytes. (B) Scatter plot of the CTCF values of the intensity of ER-Tracker [p<0.001; HC vs PD] (Data are mean ± SD, Two Sample t-Test; n=50). (C-D) qPCR. (C-D) Quantification of mRNA gene expression of GADD34 (C) and CHOP (D) in HC and PD astrocytes using quantitative polymerase chain reaction (qPCR) [p<0.001; HC vs PD] (Data are mean ± SD, Two Sample t-Test; n=50). (E-F) Graphical representation of the flow cytometry quantified positive population for p-PERK (E) and CHOP (F) in HC and PD astrocytes [p<0.001; HC vs PD] (Data are mean ± SD, Two Sample t-Test; n=5). (G-H) Graphical representation of the flow cytometry quantified positive population for LAMP1 (G) and LAMP2 (H) in EV and IV transfected U87 cells [p<0.001; EV vs IV] (Data are mean ± SD, Two Sample t-Test; n=5). (I) Representative Click-iT™ HPG Alexa Fluor™ 488 images show newly synthesized proteins (green) with nuclei counterstained with DAPI in HC and PD astrocytes. (J) Scatter plot of the CTCF values of the intensity of Click-iT™ HPG Alexa Fluor™ 488 [p<0.001; HC vs PD] (Data are mean ± SD, Two Sample t-Test; n=50).

**Figure 8.**
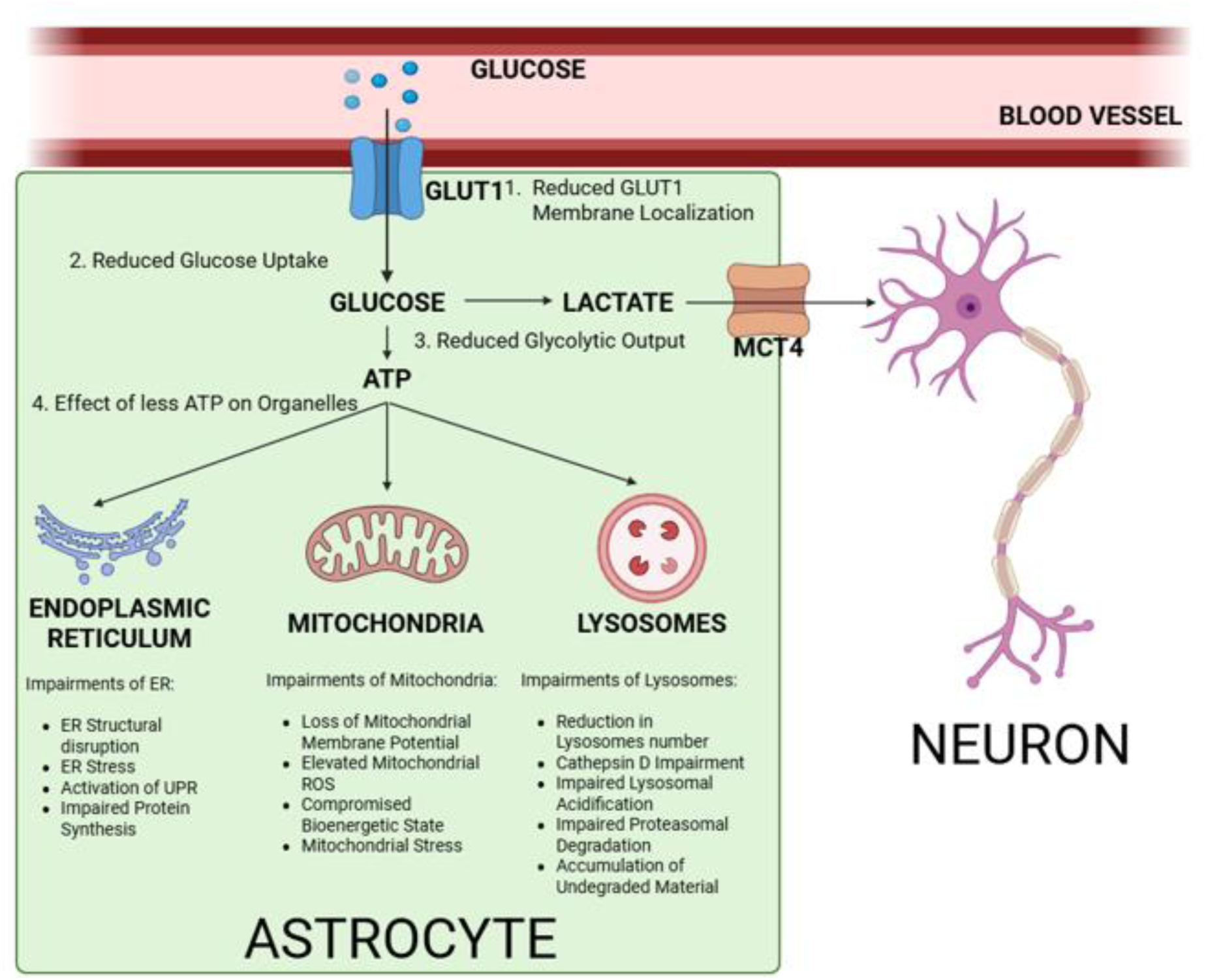
Schematic Representation Depicting the Impact of the LRRK2 I1371V Mutation on Astrocytic Energy Metabolism and Organelle Function. The LRRK2 I1371V mutation reduces GLUT1-mediated glucose uptake and ATP generation, contributing to ER stress, mitochondrial dysfunction, and lysosomal impairment, collectively weakening neuronal support.

## Discussion

The brain runs almost exclusively on glucose. Astrocytes orchestrate this energy supply through GLUT1 transporters at their end-foot processes, taking up glucose, converting it to lactate via glycolysis, and shuttling this lactate to neurons through MCT4 transporters (Gonçalves et al., 2019). Break this shuttle and neurons starve. Disruption of the astrocyte-neuron lactate shuttle drives neurodegeneration in PD (Cheng et al., 2021; Singh et al., 2025), where compromised astrocytic lactate supply leaves neurons vulnerable to energy depletion and oxidative stress. Both astrocytic glycolysis and overall brain glucose metabolism decline with age and in neurodegenerative conditions (Beard et al., 2022). LRRK2 mutations influence glucose regulation and insulin signalling beyond their known brain effects (Funk et al., 2018; Imai et al., 2020; Tsafaras et al., 2022; Kawakami et al., 2023). Impaired glucose handling, reduced oxidative phosphorylation, and depleted ATP levels drive the energy crisis in PD (Saxena, 2012; Cunnane et al., 2020). Any reduction in astrocytic lactate production restricts neuronal energy supply and amplifies vulnerability to stress.

We report the first comprehensive analysis of glucose metabolism and organellar function in LRRK2 I1371V astrocytes. I1371V astrocytes exhibit a previously unrecognized metabolic vulnerability: impaired glucose uptake driven by reduced GLUT1 expression and defective membrane localization, culminating in decreased lactate production. This primary metabolic deficit triggers cascading dysfunction across mitochondria, lysosomes, and the endoplasmic reticulum. These findings establish how LRRK2 GTPase domain mutations drive astrocytic energy failure and identify mutation-specific pathogenic mechanisms in LRRK2-associated PD.

Patient-derived astrocytes showed significantly reduced GLUT1 expression, both overall and at the plasma membrane—an effect independently confirmed in U87 cells expressing the I1371V variant. Compared with G2019S, the contrast is striking: iPSC-derived astrocytes carrying this kinase domain mutation show no GLUT1 mRNA deficits or glucose uptake impairment (Sonninen et al., 2020), and ATP synthesis remains normal (Di Domenico et al., 2019). This diverges sharply from the reduced ATP production we documented in I1371V astrocytes (Banerjee et al., 2023). The implication is clear: distinct LRRK2 mutations differentially impact astrocytic energy metabolism, with I1371V creating a unique metabolic vulnerability. The decreased 2-NBDG uptake observed in PD astrocytes reflects a substantial impairment in glucose transport and metabolic utilization. Since 2-NBDG, a non-metabolizable fluorescent analogue of glucose and it enters cells primarily via GLUT transporters, the reduction in fluorescence intensity directly indicates diminished glucose uptake capacity. Functionally, this led to a marked decrease in lactate release, indicating impaired glycolytic flux. The concomitant downregulation of LDH transcripts and the lactate exporter MCT4 further points to a collapse in the astrocytic metabolic pathway that normally couples glycolysis to neuronal energy support. Impairment of this shuttle could deprive neurons of a critical energy substrate (Yang et al., 2023). Thus, exposing them to excitotoxic and oxidative stress (Dai et al., 2023), both of which are central features of PD pathology. Importantly, in our earlier work (Banerjee et al., 2023), we also demonstrated that the I1371V mutation disrupts Nrf2 mediated glutathione metabolism, further weakening neuronal antioxidant defences.

What drives the reduced GLUT1 expression and membrane localization? Multiple converging pathways are at play. LRRK2 regulates vesicular trafficking through phosphorylation of Rab GTPases, particularly Rab8A and Rab10, which control membrane sterol and protein trafficking and recycling (Steger et al., 2016; Lis et al., 2018; Potdar et al., 2024). Hyperphosphorylation of these Rabs disrupts normal recycling and delivery of membrane cholesterol and proteins to the cell surface (Rivero-Ríos et al. 2019), compromising both membrane composition and proper localization of critical transporters and receptors (Lazarevic et al. 2022; Morissette et al. 2018). The downstream effects of aberrant Rab GTPase activity extend to alterations in membrane fluidity, ultimately impairing the functional presentation of proteins at the plasma membrane. Our previous work demonstrated hyperphosphorylation of Rab8A and Rab10 in I1371V astrocytes (Banerjee et al., 2023) and in Floor Plate Cells affecting membrane cholesterol docking, membrane fluidity and cell surface docking of receptors (Potdar et al., 2024). Recent studies have also shown that LRRK2 phosphorylates Rab8A in a glucose-dependent manner to regulate ciliary trafficking (Dhekne et al., 2018). Given that Rab8A regulates GLUT4 translocation in adipocytes (Kawakami et al., 2023), it is plausible that aberrant Rab8A phosphorylation in I1371V astrocytes disrupts GLUT1 trafficking to the plasma membrane, thereby reducing glucose uptake capacity. This interpretation is consistent with our observation of reduced cell-surface GLUT1, despite residual total cellular expression, suggesting impaired transporter trafficking rather than complete loss of synthesis.

The severe mitochondrial impairment in I1371V astrocytes—reduced membrane potential, elevated reactive oxygen species, and increased mitochondrial ubiquitination—follows directly from glucose deprivation. Mitochondrial homeostasis depends absolutely on adequate energy supply (McNair et al., 2019 & 2020; Zhuo et al., 2016; Malmgren et al., 2009). Maintaining mitochondrial membrane potential requires constant ATP input to generate ion gradients, and glucose limitation directly compromises this process (Halestrap, 1987; Nicholls, 2004). In our study, astrocytes carrying the LRRK2 I1371V mutation showed a significant reduction in JC-1 red-to-green fluorescence ratio, indicating impaired oxidative phosphorylation (Qadri et al., 2018). We saw this effect consistently across both iPSC-derived astrocytes and U87 cells expressing the I1371V variant, underscoring the mutation’s direct contribution to organellar dysfunction and reduced ATP synthesis. Similar defects have been described in other pathogenic LRRK2 variants, including G2019S (Pereira et al., 2014; Howlett et al., 2017; Cooper et al., 2012; Sanders et al., 2014), R1441G (Bahnassawy et al., 2013; Singh et al., 2019), and R1441C (Wang et al., 2012).

The elevated mitochondrial ROS observed here might be due to the compromised NADPH substrate required for regulating mitochondrial ROS. NADPH is generated through the Pentose Phosphate Pathway, which is dependent on glucose metabolism (Dikalov et al. 2011;Ge et al. 2020).

Although dysfunctional mitochondria in LRRK2 I1371V PD astrocytes were appropriately tagged for removal, as evidenced by enhanced Ubiquitin, VDAC1 colocalization, reduced 20S proteasomal activity would have resulted in inefficient clearance; and proteosomal activity known to be highly dependent on ATP (de la Peña et al. 2018). Negligible alterations in mitochondrial mass and fission-fusion balance indicate that the I1371V mutation induces an early, functionally stressed mitochondrial state rather than overt structural damage. Mitochondrial dysfunction in LRRK2 models is well documented across multiple mutations (Mortiboys et al., 2010; Yue et al., 2015; Bonello et al., 2019; Walter et al., 2019), but most studies examine these defects in isolation from metabolic context. We show that glucose uptake and metabolism deficits occur concurrently with mitochondrial impairment in LRRK2 I1371V astrocytes, suggesting that therapeutic strategies that bypass glucose uptake deficits—such as ketone body supplementation or direct lactate administration—could ameliorate mitochondrial dysfunction and improve cellular bioenergetics.

ATP depletion in LRRK2 I1371V astrocytes extends far beyond energy shortages—it dismantles cellular quality control. The comprehensive lysosomal deficits identified in I1371V astrocytes—encompassing reduced LAMP1/LAMP2 expression, impaired acidification, diminished Cathepsin D expression and activity, and accumulation of undegraded cargo—represent a critical failure in cellular proteostasis with direct relevance to PD pathogenesis. Lysosomes serve as the terminal degradative compartment for autophagy, and their dysfunction links directly to α-synuclein accumulation and PD progression (Dehay et al., 2013; Mazzulli et al., 2016). That Cathepsin D specifically is compromised while Cathepsins B and L remain functional is particularly noteworthy—Cathepsin D is the primary protease responsible for α-synuclein degradation in lysosomes (Cullen et al., 2009; Qiao et al., 2008). Mutations in Cathepsin D cause neuronal ceroid lipofuscinosis and Parkinsonism (Cullen et al., 2009; Chiu et al., 2024; Siintola et al., 2006; Koch et al., 2011), and reduced Cathepsin D activity appears in the brains of sporadic PD patients (Papagiannakis et al., 2019). Even mutations in the GBA1 gene in neurons increase monomeric α-synuclein levels by affecting lysosomal cathepsin D (Yang et al., 2020).

The impaired lysosomal acidification observed by LysoSensor imaging mechanistically explains the reduced Cathepsin D activity. This aspartic protease requires acidic pH (3.5-5.0) for maximal function (Barrett, 1977). Maintaining lysosomal acidity demands substantial ATP from vacuolar H+-ATPases (v-ATPases), making acidification exquisitely sensitive to cellular energy status (Collins et al., 2020; Zeng et al., 2020; Mindell, 2012). The glucose uptake deficits and reduced ATP production in LRRK2 I1371V astrocytes directly compromise v-ATPase function, alkalinizing the lysosomal lumen and inactivating pH-dependent proteases.

Fewer, less acidic lysosomes create a bottleneck in cargo degradation (Lee et al., 2011), thereby explaining the accumulation of undegraded vesicular material upon α-synuclein exposure. Transmission electron microscopy reveals lysosomes packed with undegraded vesicular material in LRRK2 I1371V astrocytes following α-synuclein exposure—compelling ultrastructural evidence of compromised degradative capacity. This observation has critical implications for understanding how LRRK2 I1371V astrocytes might contribute to disease progression: if astrocytes cannot efficiently degrade internalised α-synuclein, they may become sources of pathological protein propagation rather than protective clearance mechanisms. Previous studies have demonstrated that astrocytes can internalise neuron-derived α-synuclein and either degrade it or re-release it in their secretome (Raj et al., 2024; Lee et al., 2012; Rostami et al., 2017). LRRK2 I1371V astrocytes default toward the latter pathway due to lysosomal insufficiency, potentially facilitating cell-to-cell transmission of pathology.

Given how energy-intensive lysosomal biogenesis, acidification, and proteolytic activity are, there’s a clear mechanistic link between the glucose uptake deficits and lysosomal dysfunction we observed. This connection is supported by studies showing that metabolic stress and ATP depletion compromise lysosomal function and autophagy flux (Jeon et al., 2016; Raben and Puertollano, 2016). The transcription factor EB (TFEB), master regulator of lysosomal biogenesis, requires energy-dependent nuclear translocation and is inhibited under energy stress (Settembre et al., 2012; Roczniak-Ferguson et al. 2012), explaining the reduced LAMP1/LAMP2 expression in metabolically compromised LRRK2 I1371V astrocytes. ATP shortage impairs autophagy—an inherently energy-demanding process (Li et al., 2017). Autophagosome formation, vesicle transport, and fusion all require ATP. Motor proteins such as dynein and kinesin require ATP to position lysosomes and facilitate their fusion with autophagosomes. Compromised mitochondria produce excess ROS, which peroxidizes lysosomal membranes, causing lysosomal membrane permeabilization and leakage of hydrolases into the cytosol—events that trigger inflammatory signaling (He et al. 2023). We found pro-inflammatory cytokines IL6, IL1β, and TNFα released by PD astrocytes were significantly elevated in our previous study.

Energy deficits also impacted ER function, a central hub for other energy intensive cellular processes, most notably protein synthesis. ATP is critical for multiple aspects of protein translation, including ribosome function, tRNA charging, peptide elongation, and proper folding of nascent polypeptides within the ER (Thomson et al., 2013; Ibba et al., 2000; Chaudhry et al., 2003). ER Tracker staining revealed a significant reduction in ER signal in PD astrocytes, indicating structural disruption and elevated ER stress. Activation of ER stress pathways in LRRK2 I1371V astrocytes (upregulation of GADD34, CHOP, and phosphorylated PERK) coupled with reduced nascent protein synthesis reflects the cellular response to energy insufficiency. The ER is responsible for folding approximately one-third of cellular proteins and maintaining calcium homeostasis, both highly ATP-dependent processes (Schröder and Kaufman, 2005; Kozutsumi et al., 1988). Energy depletion compromises the ER folding machinery, causing accumulation of unfolded proteins and activation of the unfolded protein response (UPR). Transcriptional upregulation of UPR markers GADD34 and CHOP and increased levels of p-PERK and CHOP at the protein level confirm ER stress response activation. Chronic ER stress inhibits global protein synthesis—the Click-iT assay showed significantly reduced nascent protein production in PD astrocytes. ER stress drives PD pathogenesis, with evidence from both patient tissue and experimental models (Hoozemans et al., 2007; Mercado et al., 2016). LRRK2 G2019S studies report ER stress activation through SERCA activity modulation and enhanced sensitivity to α-synuclein-induced ER dysfunction (Lee et al., 2019; Yao et al., 2020). Our findings extend these observations to the LRRK2 I1371V mutation and position metabolic insufficiency as an upstream driver of ER stress, not merely a consequence of protein misfolding.

Reduced protein synthesis capacity in LRRK2 I1371V astrocytes has profound implications for neuronal support. Astrocytes synthesize and secrete numerous neurotrophic factors, antioxidant proteins, and metabolic enzymes critical for neuronal health (Ganapathy et al, 2018&2019; Allen and Eroglu, 2017). Impaired protein synthesis reduces production of these supportive factors, including glial cell-derived neurotrophic factor (GDNF), brain-derived neurotrophic factor (BDNF), and glutathione—all particularly important for dopaminergic neuron survival (Ganapathy et al, 2018&2019; Banerjee et al., 2023; Airavaara et al., 2012; Dringen et al., 2000). This proteostatic failure represents another mechanism through which LRRK2 I1371V astrocytes fail to provide adequate neurotrophic support.

Our findings support a unified model: glucose uptake deficits in LRRK2 I1371V astrocytes trigger organellar dysfunction that propagates through the integrated mitochondria-ER-lysosome axis. Organellar crosstalk networks maintain cellular equilibrium—dysfunction in one compartment reverberates throughout interconnected organelles via shared metabolic pathways, calcium signalling, and membrane contact sites (Rieusset, 2018; Krols et al., 2016).

In LRRK2 I1371V astrocytes, reduced glucose uptake and ATP depletion compromise maintenance of mitochondrial membrane potential, thereby increasing ROS production. Elevated ROS damage mitochondrial DNA and proteins, thereby impairing respiratory chain function and creating a feed-forward cycle of oxidative stress (Kowaltowski et al., 2009). Meanwhile, ATP depletion compromises lysosomal acidification through impaired v-ATPase function, reducing proteolytic capacity and causing accumulation of damaged organelles and protein aggregates. The ER—deprived of ATP for chaperone function and bombarded by oxidative stress from dysfunctional mitochondria—activates the UPR and shuts down protein synthesis, further compromising cellular function.

This integrated dysfunction explains why therapeutic interventions targeting individual organelles in isolation have failed in PD models (Snow et al. 2010, Udayar et al. 2022). Restoring glucose metabolism and energy homeostasis represents a more fundamental therapeutic approach—one that normalizes function across multiple organellar compartments simultaneously. Metabolic therapies for neurodegenerative diseases—ketogenic diets, exercise interventions, and pharmacological metabolic modulators—are attracting increased attention for precisely this reason (Norwitz et al., 2019; Bohnen et al., 2022).

Our findings point to therapeutic strategies that move past traditional dopamine replacement. Astrocytic glucose metabolism appears as a novel therapeutic target in LRRK2 I1371V-associated PD. Approaches to improve glucose uptake or bypass impaired glucose metabolism could restore astrocytic support function and protect dopaminergic neurons. Metabolic substrate supplementation offers one avenue: administering alternative energy substrates, such as ketone bodies (β-hydroxybutyrate), medium-chain triglycerides, or lactate, bypasses glucose uptake deficits and directly fuels oxidative metabolism. Ketogenic diets show promise in preclinical PD models (Vanitallie et al., 2005; Yang and Cheng, 2010), and our findings suggest a particular benefit for LRRK2 I1371V patients by compensating for deficits in astrocytic glucose metabolism. GLUT1 upregulation provides another approach. Therapeutic strategies that increase GLUT1 expression or improve its trafficking to the plasma membrane restore glucose uptake capacity, possibly via activation of transcriptional regulators such as hypoxia-inducible factor 1α (HIF-1α) or modulation of trafficking machinery. Certain compounds enhance GLUT1 expression in brain endothelial cells, suggesting feasibility (Patching, 2017). Rab GTPase modulation addresses the root cause. Given aberrant Rab8A/Rab10 phosphorylation in GLUT1 mislocalization, normalizing Rab function could restore glucose transporter trafficking. LRRK2 kinase inhibitors, currently in clinical development for PD (Dzamko et al., 2017; Tolosa et al., 2020), might achieve this effect, though their efficacy in LRRK2 I1371V patients requires investigation since this mutation mainly impacts GTPase rather than kinase function. Enhancing lysosomal function offers yet another strategy. Compounds that improve lysosomal acidification or enhance Cathepsin D activity restore protein clearance capacity even in metabolically stressed astrocytes. Glucocerebrosidase modulators, currently under development based on the GBA-PD connection (Migdalska-Richards and Schapira, 2016), might benefit lysosomal function across multiple PD etiologies. Finally, targeted antioxidants—particularly those designed to accumulate in mitochondria such as MitoQ or SS-31—could ameliorate oxidative damage and improve cellular bioenergetics given the elevated ROS and mitochondrial dysfunction in I1371V astrocytes (Snow et al., 2010; Ghosh et al., 2010).

Metabolic interventions are widely accepted and can be initiated in at-risk individuals (LRRK2 mutation carriers) before motor symptoms appear, potentially postponing or even preventing disease progression. This aligns with the broader trend toward early initiation of disease-modifying therapies in the neurodegenerative process (Simon et al., 2020; Feigin et al., 2017).

Comparative studies examining other Roc-COR domain mutations (R1441C/G/H, N1437H, Y1699C) will determine whether glucose metabolism deficits are shared across GTPase domain mutations or specific to I1371V. Longitudinal studies examining when metabolic deficits emerge during disease progression—ideally in presymptomatic I1371V carriers—will identify windows for preventive intervention. Developing biomarkers that reflect astrocytic metabolic dysfunction (through PET imaging of glucose metabolism or CSF metabolomic profiling) will enable patient stratification and monitoring of therapeutic responses.

## Conclusion

The LRRK2 I1371V mutation disrupts energy balance in astrocytes through a connected cascade of events. Reduced GLUT1 mediated glucose uptake limits glycolysis and decreases lactate supply to neurons. Mitochondrial dysfunction, marked by depolarization and higher ROS, worsens this energy stress. Proteasomal activity is compromised, and lysosomes show deficits such as lower LAMP1 and LAMP2, impaired acidification, and selective loss of Cathepsin D, which hinders clearance of damaged organelles. ER stress and activation of the unfolded protein response suppress overall protein synthesis in turn manifested by lower levels of lysosomal enzymes and transporters such as GLUT1 and SLC1A2. These deficits form a vicious cycle of energy depletion, impaired proteostasis, and organelle dysfunction feed into each other. These insights open new avenues for therapeutic development focused on restoring astrocytic metabolic capacity and support functions, potentially offering disease-modifying benefits when initiated early in the disease course.

## Data availability

All data generated or analysed during this study are included in this published article [and its supplementary information files].

## Acknowledgements

We acknowledge Dr Manjunath, Department of Neurovirology, NIMHANS, for access to the Advanced Flow Cytometer facility and Dr P. Govindaraj, Department of Neuropathology, NIMHANS, for access to the TEM facility. We acknowledge financial support from a grant from the Department of Biotechnology (DBT), Government of India, New Delhi; contract grant No. BT/PR45527/MED/122/322/2022 by ID and Parkinson’s Disease & Movement Disorder Research Fund (PDMDRF), NIMHANS by VH, NK, RY, PKP. RB is supported by UGC NET-JRF Ph.D.

## Author Contribution Statement

Conceptualization: I.D; methodology: R.B, R.S and I.D; formal analysis and investigation: R.B, I.D, R.S, R.Y, P.K.P, V.H and N.K; writing - original draft preparation: R.B and I.D; writing-review and editing: I.D, R.B; R.S, R.Y, P.K.P, V.H and N.K; funding acquisition: I.D, R.Y, and P.K.P, V.H and N.K; resources: I.D, R.Y, P.K.P, V.H and N.K; supervision: I.D. All authors read and approved the final manuscript.

---

**Fig. S1.**
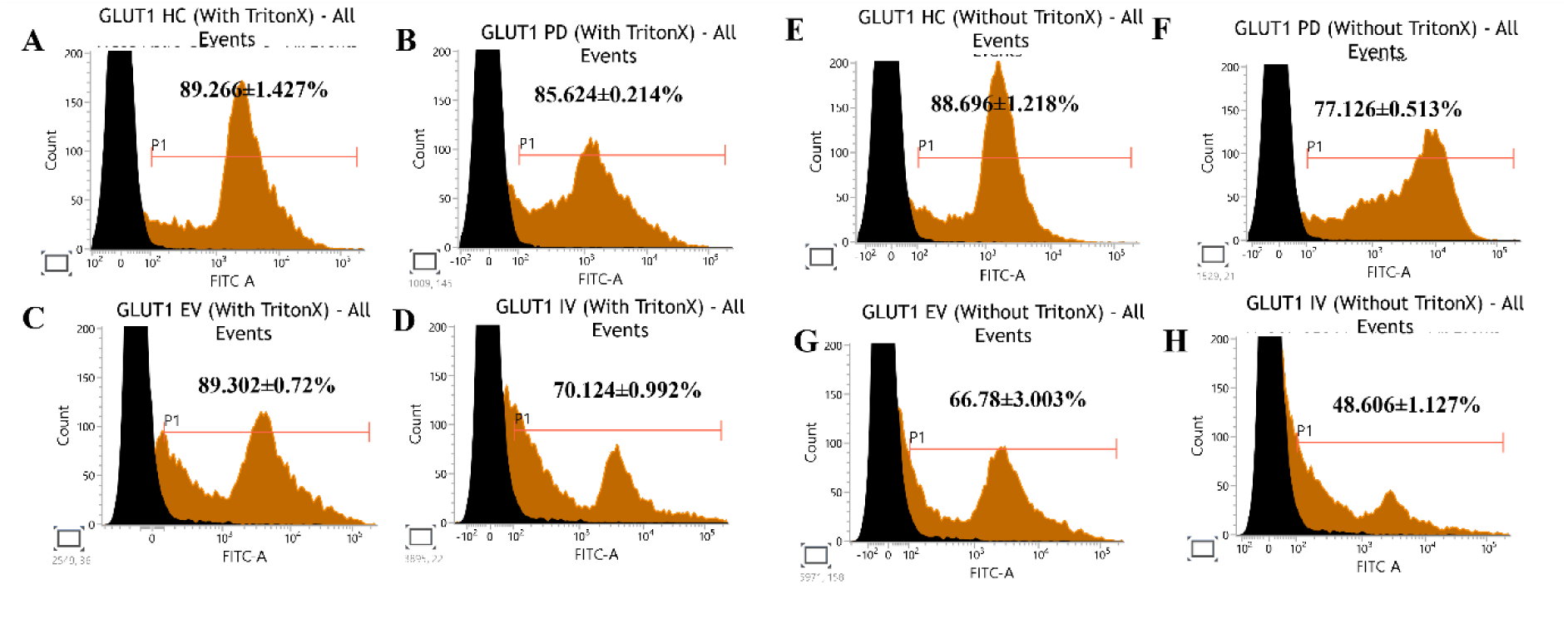
**(A-B)** Representative flow cytometry histogram of total GLUT1 in HC **(A)** and PD **(B)** astrocytes. **(C-D)** Representative flow cytometry histogram of total GLUT1 in EV **(C)** and IV **(D)** transfected U87 cells. **(E-F)** Representative flow cytometry histogram of membrane localized GLUT1 in HC **(E)** and PD **(F)** astrocytes. **(G-H)** Representative flow cytometry histogram of total GLUT1 in EV **(G)** and IV **(H)** transfected U87 cells.

**Fig. S2.**
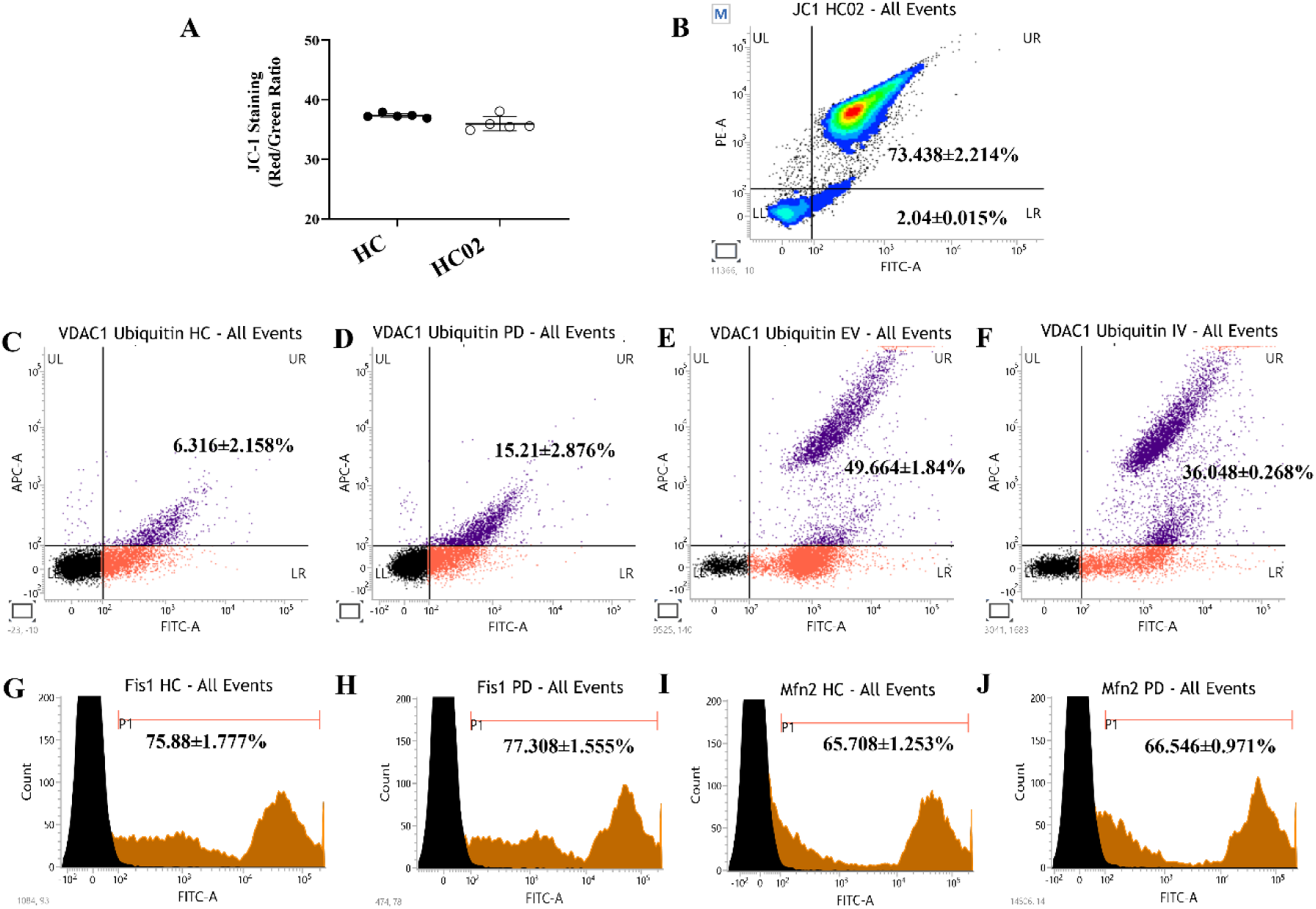
**(A)** Quantification of JC-1 aggregate/monomer ratios in HC and HC02 Astrocytes. (n=5) **(B)** Representative JC-1 flow cytometry histogram illustrating red (aggregate) to green (monomer) fluorescence in HC02 astrocytes. **(C-D)** Representative Scatter Plot of dual positive VDAC1 and Ubiquitin population of cells in HC **(C)** and PD **(D)** astrocytes. **(E-F)** Representative Scatter Plot of dual positive VDAC1 and Ubiquitin population of cells in EV **(E)** and IV **(F)** transfected U87 cells. **(G-H)** Representative flow cytometry histogram of LAMP1 in HC **(G)** and PD **(H)** astrocytes. **(I-J)** Representative flow cytometry histogram of LAMP2 in HC **(I)** and PD **(J)** astrocytes.

**Fig. S3.**
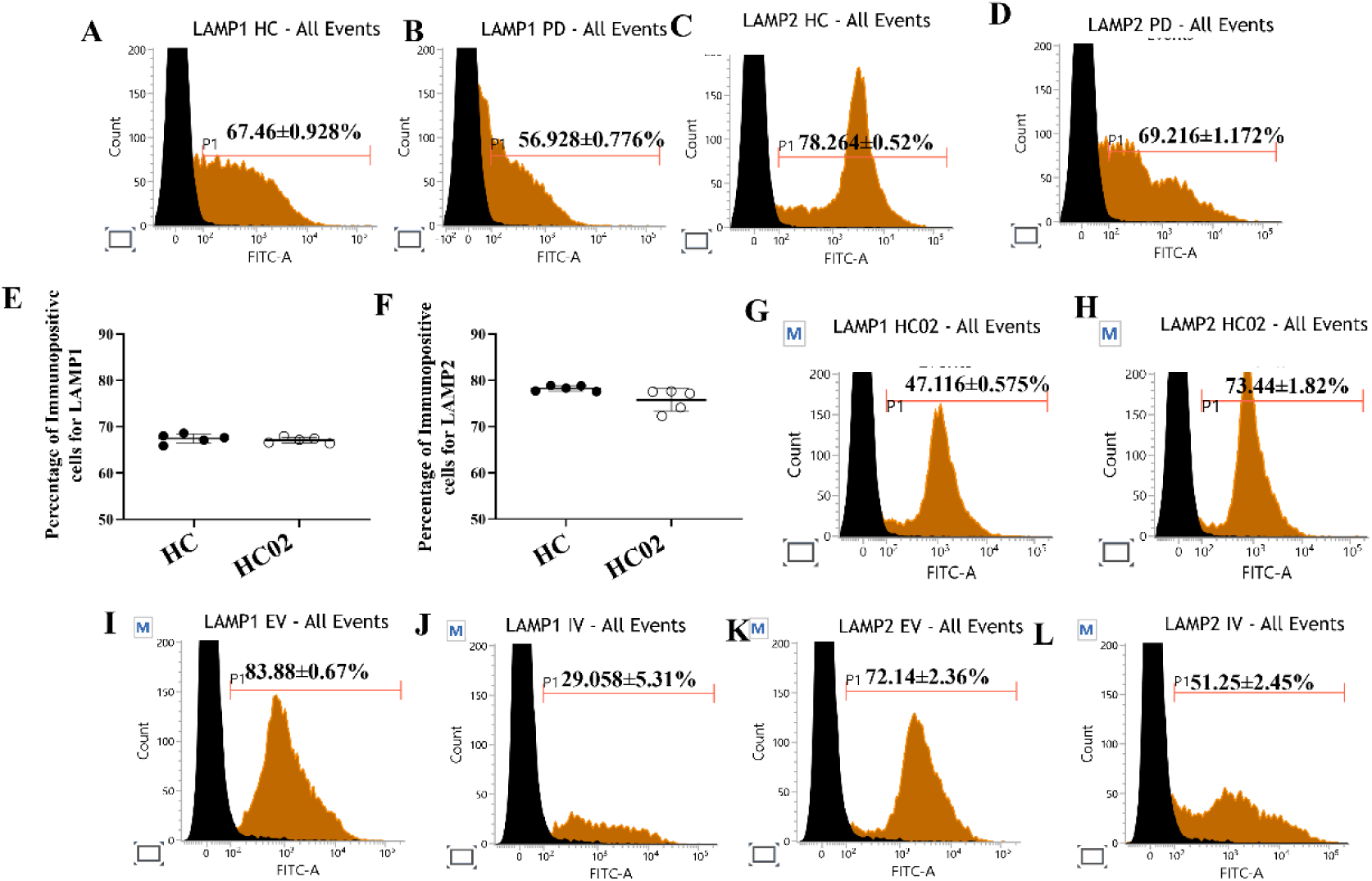
**(A-B)** Representative flow cytometry histogram of LAMP1 in HC **(A)** and PD **(B)** astrocytes. **(C-D)** Representative flow cytometry histogram of LAMP2 in HC **(C)** and PD **(D)** astrocytes. **(E)** Graphical representation of the flow cytometry quantified positive population for LAMP1 in HC and HC02 astrocytes. (n=5) **(F)** Graphical representation of the flow cytometry quantified positive population for LAMP2 in HC and HC02 astrocytes. (n=5) **(G)** Representative flow cytometry histogram of LAMP1 in HC02 astrocytes. **(H)** Representative flow cytometry histogram of LAMP2 in HC02 astrocytes. **(I-J)** Representative flow cytometry histogram of LAMP1 in EV **(I)** and IV **(J)** transfected U87 cells. **(K-L)** Representative flow cytometry histogram of LAMP2 in EV **(K)** and IV **(L)** transfected U87 cells.

**Fig. S4.**
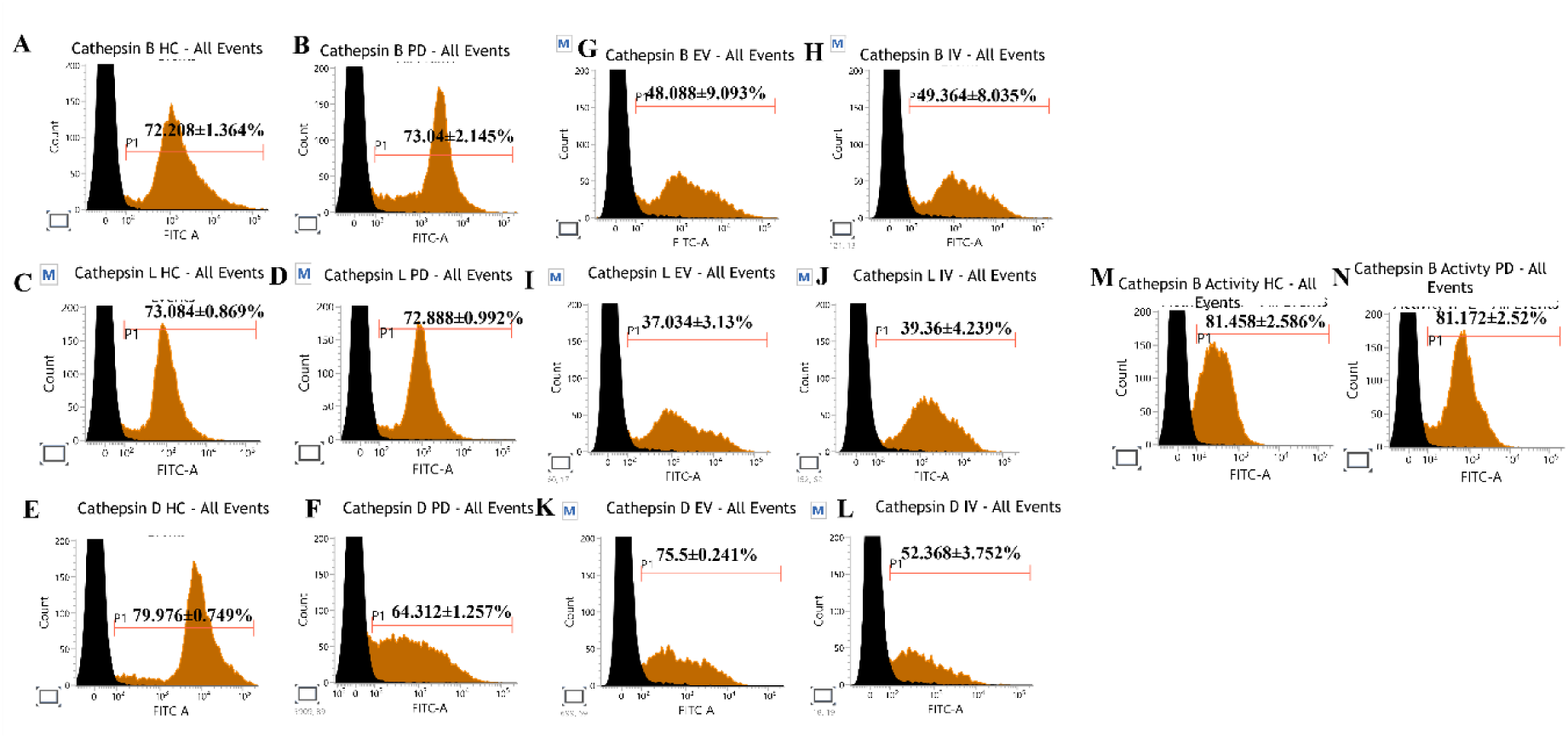
Flow Cytometry Histograms. **(A-B)** Representative flow cytometry histogram of Cathepsin B in HC **(A)** and PD **(B)** astrocytes. **(C-D)** Representative flow cytometry histogram of Cathepsin L in HC **(C)** and PD **(D)** astrocytes. **(E-F)** Representative flow cytometry histogram of Cathepsin D in HC **(E)** and PD **(F)** astrocytes. **(G-H)** Representative flow cytometry histogram of Cathepsin B in EV **(G)** and IV **(H)** transfected U87 cells. **(I-J)** Representative flow cytometry histogram of Cathepsin L in EV **(I)** and IV **(J)** transfected U87 cells. **(K-L)** Representative flow cytometry histogram of Cathepsin D in EV **(K)** and IV **(L)** transfected U87 cells. **(M-N)** Representative flow cytometry histogram of Cathepsin D in HC **(M)** and PD **(N)** astrocytes.

**Fig. S5.**
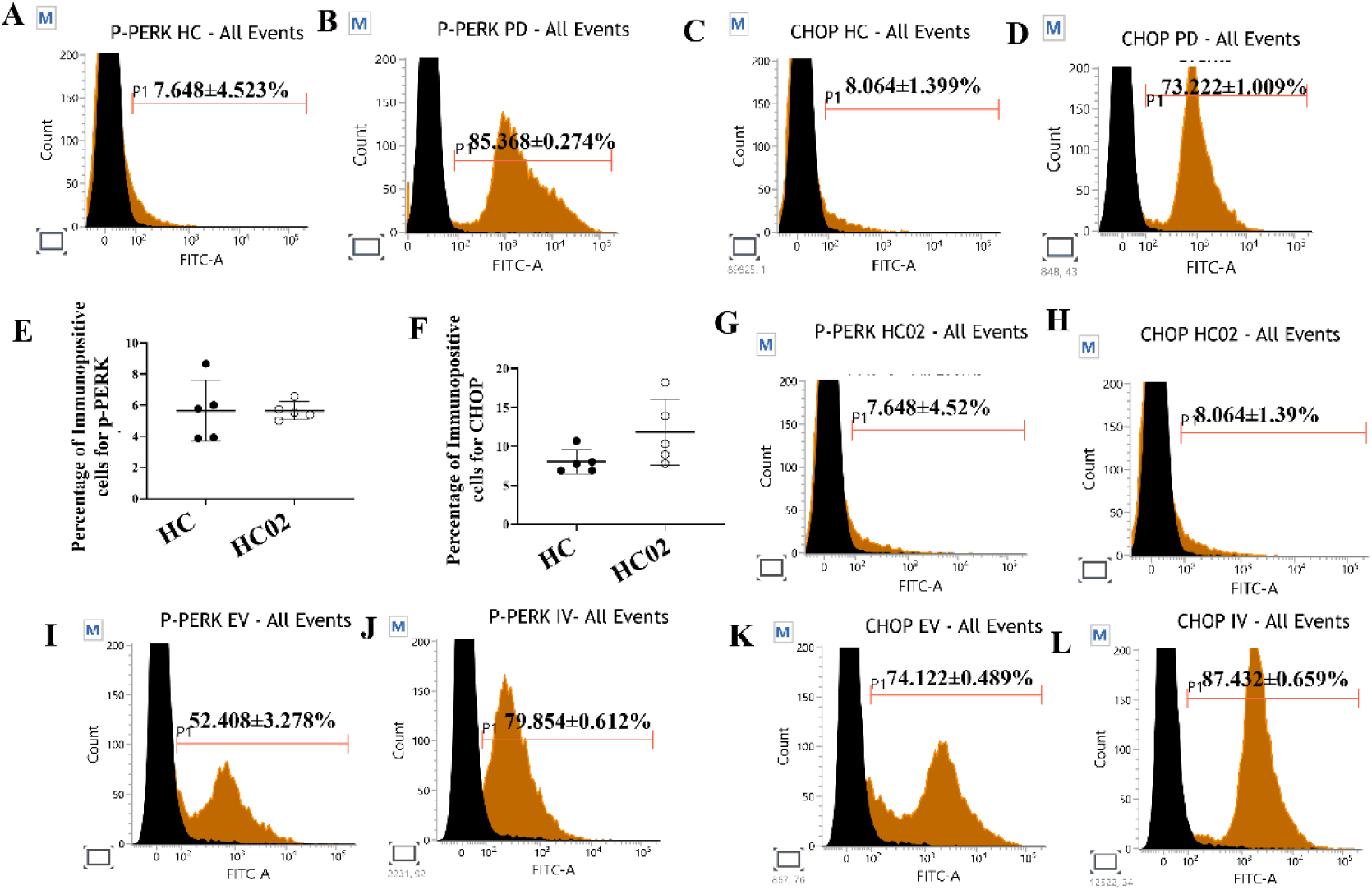
Flow Cytometry Histograms. **(A-B)** Representative flow cytometry histogram of p-PERK in HC **(A)** and PD **(B)** astrocytes. **(C-D)** Representative flow cytometry histogram of CHOP in HC **(C)** and PD **(D)** astrocytes. **(E)** Graphical representation of the flow cytometry quantified positive population for p-PERK in HC and HC02 astrocytes. (n=5) **(F)** Graphical representation of the flow cytometry quantified positive population for CHOP in HC and HC02 astrocytes. (n=5). **(G)** Representative flow cytometry histogram of p-PERK in HC02 astrocytes. **(H)** Representative flow cytometry histogram of CHOP in HC02 astrocytes. **(I-J)** Representative flow cytometry histogram of p-PERK in EV **(I)** and IV **(J)** transfected U87 cells. **(K-L)** Representative flow cytometry histogram of CHOP in EV **(K)** and IV **(L)** transfected U87 cells.

